# Complete firing-rate response of neurons with complex intrinsic dynamics

**DOI:** 10.1101/028076

**Authors:** Maximilian Puelma Touzel, Fred Wolf

**Affiliations:** Department for Nonlinear Dynamics, Max Planck Institute for Dynamics and Self-organization, Goettingen, Niedersachsen, Germany; Bernstein Center for Computational Neuroscience, Goettingen, Niedersachsen, Germany; Institute for Nonlinear Dynamics, Georg-August University School of Science, Goettingen, Niedersachsen, Germany; Kavli Institute for Theoretical Physics, University of California Santa Barbara, Santa Barbara, California, United States of America

## Abstract

The response of a neuronal population over a space of inputs depends on the intrinsic properties of its constituent neurons. Two main modes of single neuron dynamics–integration and resonance–have been distinguished. While resonator cell types exist in a variety of brain areas, few models incorporate this feature and fewer have investigated its effects. To understand better how a resonator’s frequency preference emerges from its intrinsic dynamics and contributes to its local area’s population firing rate dynamics, we analyze the dynamic gain of an analytically solvable two-degree of freedom neuron model. In the Fokker-Planck approach, the dynamic gain is intractable. The alternative Gauss-Rice approach lifts the resetting of the voltage after a spike. This allows us to derive a complete expression for the dynamic gain of a resonator neuron model in terms of a cascade of filters on the input. We find six distinct response types and use them to fully characterize the routes to resonance across all values of the relevant timescales. We find that resonance arises primarily due to slow adaptation with an intrinsic frequency acting to sharpen and adjust the location of the resonant peak. We determine the parameter regions for the existence of an intrinsic frequency and for subthreshold and spiking resonance, finding all possible intersections of the three. The expressions and analysis presented here provide an account of how intrinsic neuron dynamics shape dynamic population response properties and can facilitate the construction of an exact theory of correlations and stability of population activity in networks containing populations of resonator neurons.

**Author Summary:** Dynamic gain, the amount by which features at specific frequencies in the input to a neuron are amplified or attenuated in its output spiking, is fundamental for the encoding of information by neural populations. Most studies of dynamic gain have focused on neurons without intrinsic degrees of freedom exhibiting integrator-type subthreshold dynamics. Many neuron types in the brain, however, exhibit complex subthreshold dynamics such as resonance, found for instance in cortical interneurons, stellate cells, and mitral cells. A resonator neuron has at least two degrees of freedom for which the classical Fokker-Planck approach to calculating the dynamic gain is largely intractable. Here, we lift the voltage-reset rule after a spike, allowing us to derive for the first time a complete expression of the dynamic gain of a resonator neuron model. We find the gain can exhibit only six shapes. The resonant ones have peaks that become large due to intrinsic adaptation and become sharp due to an intrinsic frequency. A resonance can nevertheless result from either property. The analysis presented here helps explain how intrinsic neuron dynamics shape population-level response properties and provides a powerful tool for developing theories of inter-neuron correlations and dynamic responses of neural populations.

## Introduction

Integration and resonance are two operational modes of the spiking dynamics of single neurons. These two modes can be distinguished from each other by observing the neuron’s signal transfer properties: how features in its input current transfer to features in its output spiking. The traditional approach to investigating neuronal transfer properties is to measure the stationary response: the time-averaged rate of firing of spikes as a function of the mean input current, or *fI-curve.* In Hodgkin’s classification [1], *Type I* membranes can fire at arbitrarily low rates, while the onset of firing in *Type II* membranes occurs only at a finite rate. This distinction arises naturally from the topology of the bifurcations that a neuron can undergo from resting to repetitive spiking [2]. In many central neurons, it is fluctuations rather than the mean input current that drive spiking, putting them in the so-called *fluctuation-driven* regime [3]. Many dynamical phenomena are nevertheless tightly linked to excitability type. For example, Type II neurons exhibit rebound spikes, subthreshold oscillations and spiking resonance (e.g. mitral cells, [4-6], respectively). The qualitative explanation for these phenomena is that the dynamical interplay of somatic conductances endow some neurons with a voltage frequency preference, i.e. a *subthreshold resonance.* This preference can contribute to a *superthreshold resonance* in the modulation of their output spiking [7]. How dynamic response properties of spiking dynamics such as resonance emerge can be directly assessed by considering the neuron’s dynamic gain.

Dynamic gain, first treated by Knight [8], quantifies the amount by which features at specific frequencies in the input current to a neuron are amplified or attenuated in its output spiking. It can accurately distinguish functional types and unveil a large diversity of phenomena shaping the response to dynamic stimuli [9-18]. Dynamic gain and response are also essential ingredients for theoretical studies of network dynamics in recurrent circuits [8,12,13,18-49]. First, they determine the stability of the population firing rate dynamics [21,25,26]. Second, they determine how input correlations between a pair of cells are transferred to output correlations [28,42,44-49], and from which self-consistent relations for correlations in recurrent circuits can be obtained.

Experimental studies have started over the past years to use dynamic gain measurements to investigate the encoding properties of cortical neuron populations [9-18]. Although theoretical studies have investigated many neuron models, very few models are known for which dynamical response can be explicitly calculated. One basic reason for this lies in the fact that Fokker-Planck equations for neuron models with two or more degrees of freedom are not solvable in general [50]. For Type II neuron models that require at least two degrees of freedom, no solvable model is known.

The simplest model capable of subthreshold resonance was introduced by Young [51] in the early theories of excitability. Later, Izhikevich formulated a structurally similar neuron [52]. Richardson and coworkers performed the first calculation of the linear response function of a neuron model capable of resonance, the Generalized Integrate-and-Fire (GIF) neuron [22,29]. Only in the limit of relatively slow intrinsic current time constant can analytical expressions for the GIF response be obtained, however. The distinct transfer properties of resonant vs. non-resonant dynamics leads to different information transfer properties. While this has been demonstrated in the mean-driven regime [53,54], no such results exist for the fluctuation-driven regime, in part due to a lack of exact analytical expressions for even the linear dynamic gain. Type II excitability and dynamic response thus are representative of the more general challenge posed by response properties of neurons with complex intrinsic dynamics.

In the current study, we derive and analyze the linear response function in the fluctuation-driven regime of a neuron model capable of resonance. We express it as a filter cascade from current to voltage to spiking. It is valid across all relevant input frequencies and over all relevant values of the intrinsic parameters. In particular, we apply to the GIF neuron model the Gauss-Rice approach in which the voltage reset after a spike is omitted. The methods generalize to additional intrinsic currents and to the full nonlinear response with spike generation. To understand how subthreshold features interact to determine a neuron’s filter characteristics, including resonance, we provide a two-dimensional representation of the response properties that completely characterizes all possible filter types. For this idealized model, we determine analytically and numerically a wide and biologically-relevant regime of validity of the derived expression.

The paper begins with the definition of the model and its numerical implementation. We then derive a general expression for the linear response in the mean channel of a Gauss-Rice neuron. In the next section, the analytical results for the response properties of the Gauss-Rice GIF neuron model are obtained. The final section then presents an analysis of the expression. For the sake of mathematical clarity, most calculations appear in the main text; the rest, including an exposition of model assumptions, are contained in the Methods.

## Results

### Definitions and methods for a population of Gauss-Rice GIF neurons

We consider the most simple hard-threshold, no-reset, GIF-type neuron capable of exhibiting resonator dynamics, whose response properties have been partially studied in [25]. A reset version of this model is treated in [29], where the population spiking response properties were calculated assuming large intrinsic time constant. In the Methods, we present a more detailed exposition of the model assumptions, and justify an additional simplification of the voltage reset after a spike. The feature that distinguishes the GIF model from the classical Leaky Integrate-and-Fire (LIF) model is that the dynamics of the voltage, *V*, is coupled to an intrinsic activity variable, *w*,

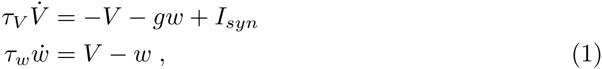

where *g* is a relative conductance and *τ_V_* and *τ_w_* are the respective time constants of the dynamics. The notation *ẋ* denotes the derivative with respect to time of the variable *x*. Spikes are emitted at upward crossings of a threshold, *θ*. Synaptic current modeled by *I_syn_* drives the model whose dynamics are kept stable by keeping *g* > −1. When *g* < 0, *w* is depolarizing. When *g* > 0, it is hyperpolarizing and can lead to resonant voltage dynamics.

#### Intrinsic parameters, *g* and *τ_w_*, shape the phase diagram of the intrinsic dynamics

Here we present analysis of the phase diagram of the intrinsic dynamics of the model, which is a reparametrization of Fig. 1 from [29]. Beyond that work, here we analyze the Ω-contour density and scaling behavior. For a fixed, constant value of *I_syn_*, and with time in units of *τ_V_*, the structure of the phase space of the single neuron dynamics described by Eq. (1) is determined by a point in the *τ_w_*/*τ_V_* vs. *g* plane, the two parameters defining the intrinsic current, *w* (see Fig. 1). For *τ_w_* ≪ *τ_V_*, *w* speeds up or slows down *V* depending on whether *g* is hyperpolarizing (*g* > 0) or depolarizing (*g* < 0) characterized by an effective time constant

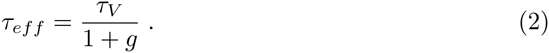

**Figure 1.**
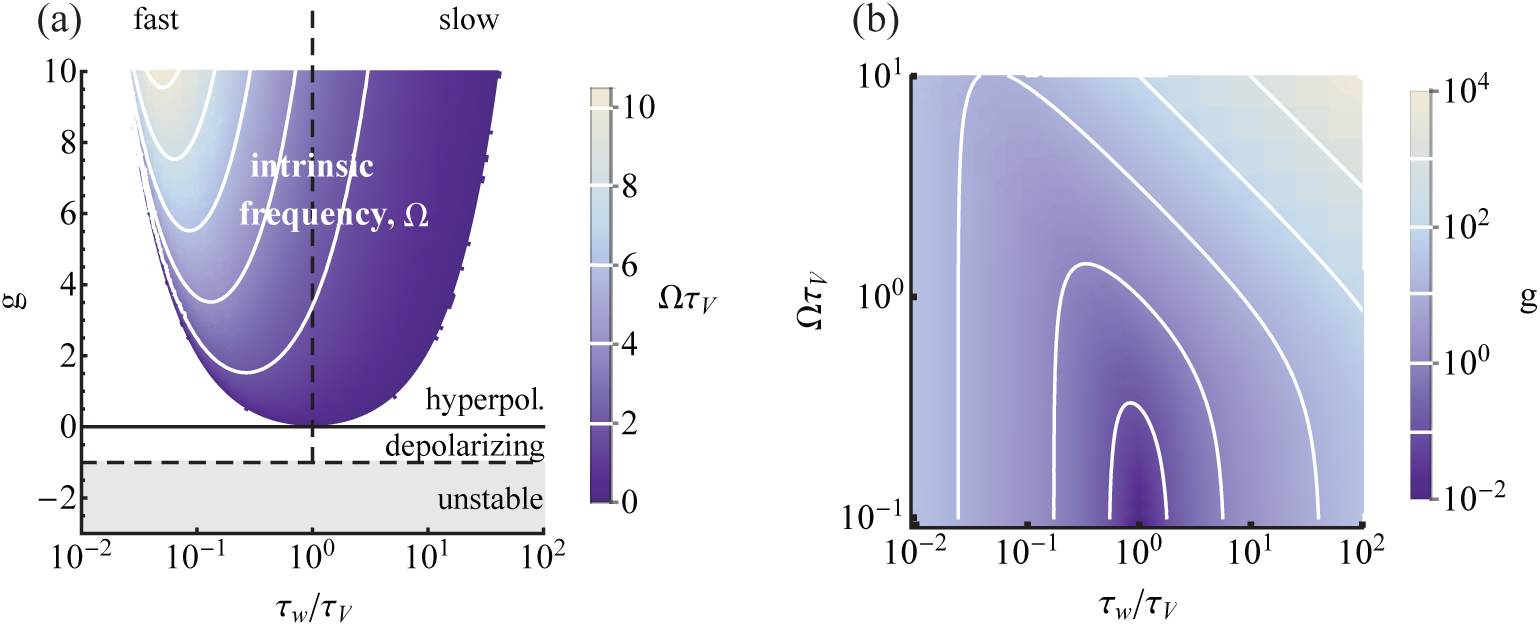
The type of *w*-current depends on the values of the intrinsic parameters. (a) Intrinsic parameter phase diagram in (*τ_w_*/*τ_V_*, *g*). *w* can be depolarizing (*g* < 0) or hyperpolarizing (*g* > 0). *w* contributes an intrinsic frequency to the model in the colored region. The dynamics are unstable if *g* < −1. Iso-Ω lines are shown in white 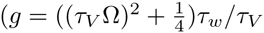 for large *τ_w_* and 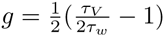 for small *τ_w_*). (b) When 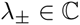, the phase diagram can be cast in (*τ_w_*/*τ_V_*, Ω*τ_V_*)-space. Iso-*g* lines are shown in white. (See [29] for a similar plot).

While the dissipative voltage term stabilizes the voltage dynamics, the dynamics can be effectively unstable for *g* < −1, and we exclude this case. For depolarizing intrinsic current, there is a region where the two eigenvalues of the voltage solution, *λ*_±_, are complex and the model exhibits an intrinsic frequency, Ω = 2*πf_int_*, that varies as

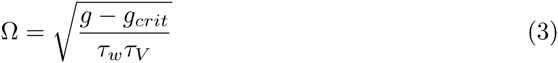

where *g_crit_* = (*τ_w_* − *τ_V_*)^2^/4*τ_w_τ_V_* (see Methods for details). For a fixed *g* > 0, a given value of Ω can be achieved at both a high and a low value of *τ_w_*. For fast *τ_w_*, the Ω-contour density is high and the model exhibits high parameter sensitivity, while for large *τ_w_* the contour density is low and the model is relatively insensitive to local parameter variation. Taking the respective limits, the set of isofrequency curves are linear for large *τ_w_* with slope ∝ Ω^2^ and 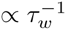 with a slope independent of Ω for small *τ_w_*.

Furthermore, there is a minimum relative conductance, 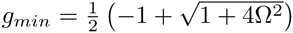 for which a given Ω can be achieved. The minimum shifts to increasing short *τ_w_* with Ω. To emphasize the timescale of the intrinsic frequency when it exists, we reparametrize the model by replacing *g* with Ω using Eq. (3), and arrive at the implicit representation shown in Table 1) (see Methods for more details). The statistical structure of the relative timings of the output spikes of the model will be affected by Ω.

**Table 1.**
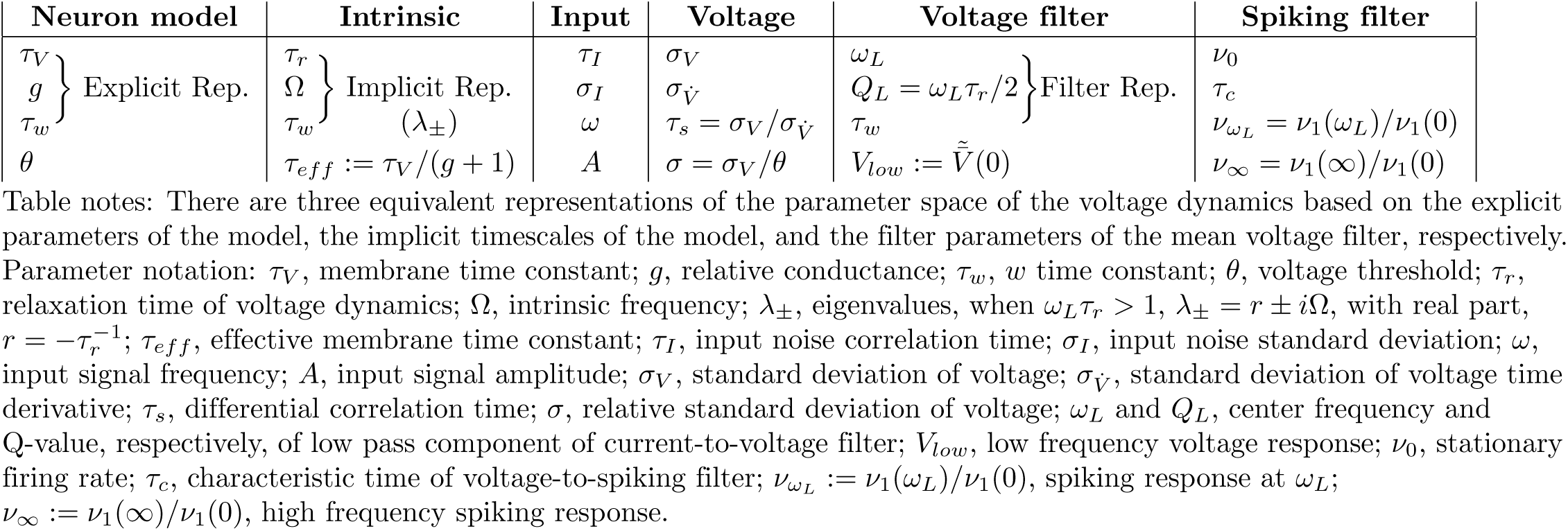
Parameter groups for each dynamics

#### Population firing rate dynamics

Given a population of *N* neurons indexed by *k*, in a time window, *T*, each one produces a spike train,

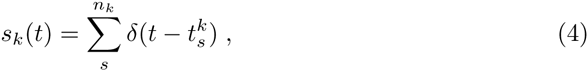

with *n_k_* spikes labeled as 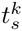. The average firing rate across the population in this window is

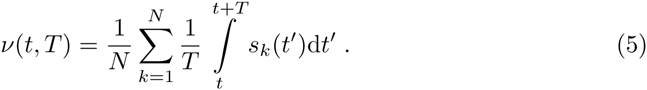

For stationary input, this becomes the stationary population averaged firing rate, independent of *t*, in the limit *T* → ∞,

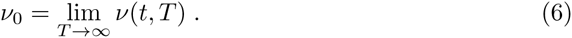

In the other limit, taking *T* → 0 while keeping *NT* constant, such that there is a statistically invariant number of spikes in the time window, the integrand of Eq. (5) is a well-defined time-dependent ensemble average, the instantaneous population firing rate,

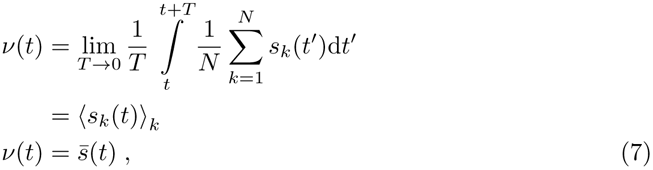

where 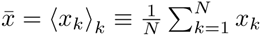 denotes the population average of a single neuron quantity, *x*. Note that this population firing rate can exhibit time dependence on arbitrarily fast timescales.

#### Populations of fluctuation-driven neurons

The input to the neuron, *I_syn_*, from Eq. (1) arrives from many, weak synapses. The total drive will thus resemble a continuous stochastic process. The system can then be solved under this assumption by directly simulating the corresponding stochastic differential system of Eq. (1),

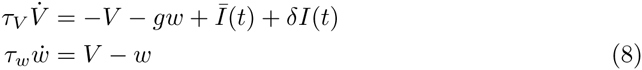

where *Ī*(*t*) is the time-dependent mean input and *δI*(*t*) is a zero-mean noise process. Solutions give the output spike times, which averaged over an ensemble give the population firing rate, *v*(*t*). Under the diffusion approximation, discussed in more detail in the Methods, the stochastic drive, *δI*(*t*), can be taken as an zero-mean Ornstein-Uhlenbeck process with variance 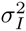 and correlation time *τ_I_*. The resulting stochastic dynamics were simulated by numerical integration via a Runge-Kutta scheme (see ref. [55] for details).

To illustrate the dynamic ensemble response, we show in Fig. 2 an example of input, intrinsic, and output variable time series produced by the model for two choices of signal in the mean channel, *Ī*(*t*): a weak oscillation of amplitude *A* and frequency *ω* and, separately, a step of height Δ. In addition, we show the corresponding population firing rate dynamics obtained from a histogram of the spike times of the sample ensemble produced by the two inputs. Code to produce this plot can be found in the supplemental material. The input modulation structures the spike times produced by the ensemble relative to the stationary response in a way that only becomes salient at this population level. We motivate consideration of the analytical expressions for the linear response function (Eq. (41)) and the step response function (Eq. (63)), obtained later in this paper, by plotting their curves, which accurately overlie the profile of the two respective measured histograms. While the input oscillation produces modulation in the output spiking at only one frequency, the step input produces a response that has power across a broad band of frequencies.

**Figure 2.**
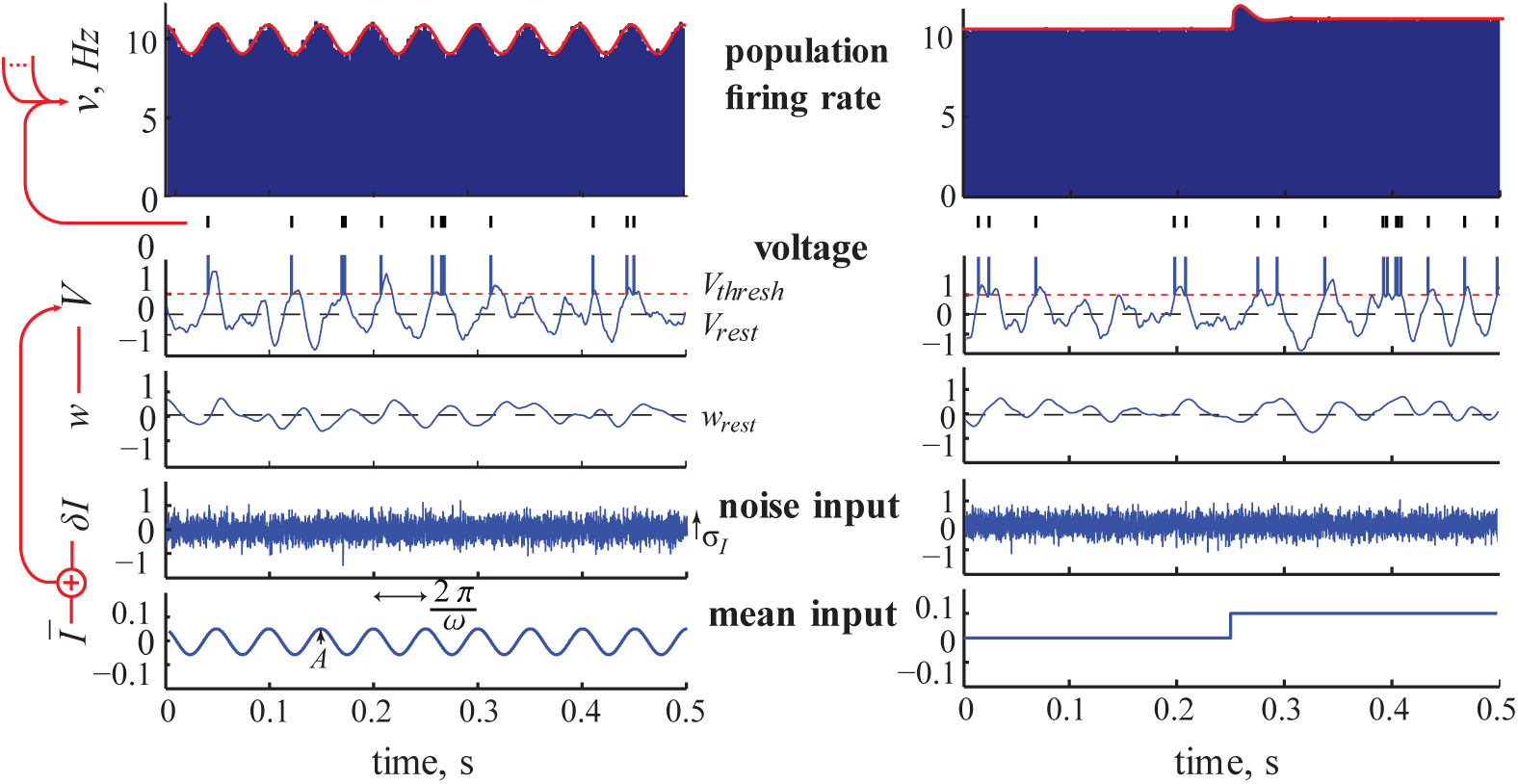
From input to ensemble response: numerics and prediction. Model output for the default parameter set: *τ_I_* = 1ms, *σ_I_* = 1, *τ_V_* = 10ms, *θ* =1, *τ_w_* = 20ms, *f_int_* = 20Hz (*g* = 3.15). Left: in in the case of an oscillation of amplitude *A* = 0.05 and input frequency *ω* = (2*π*)20 rad/s. Right: in the case of a step of height *A* = 0.1. The example realization shown is the one with the maximum number of spikes from the sample ensemble. The red line is the response calculated using the analytical expressions for the oscillation and step response, Eqs. (41) and (63), respectively.

### Approaches to obtaining the population response

Response theory captures the population response to input signals with arbitrary frequency content and so we now turn to it, and linear response theory in particular, in the pursuit of understanding the population firing rate dynamics of the GIF neuron model.

The formal, implicit definition of the linear response function, *v*_1_(*ω*), arises from a weak oscillatory modulation of amplitude *A* and frequency *ω* in the mean input, and an expansion of the response, *v*(*t*), in powers of *A*,

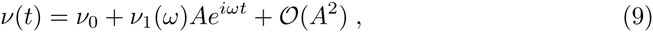

where *v*_0_ is the stationary response, Eq. (6). In the Methods, we restate how the linear response can be obtained (Eq. (59)) directly from the spike times using the complex response vector, 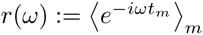. Below, we show the classic formulation that shows that it can also be obtained from the voltage dynamics.

#### Obtaining the response from the statistics of the voltage dynamics

To obtain *v*_1_(*ω*) analytically, we go back to the definition of *v*(*t*) containing *s_k_*(*t*), Eq. (7). *s_k_* (*t*) can be rewritten as

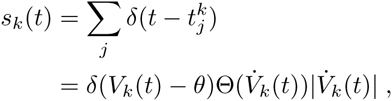

where Θ is the Heaviside theta function defined as Θ(*x*) = 0 for *x* < 1 and Θ(*x*) = 1 for *x* > 0. 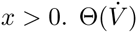 appears since spikes are only generated at *upward* threshold crossings of the voltage. The factor 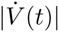 results from the coordinate change in the argument of the *δ*-function. When combined with 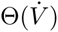, the absolute value can be omitted. For a population of such neurons, we can then obtain the population-averaged firing rate as the rate of upward threshold crossings known as Rice’s formula [56],

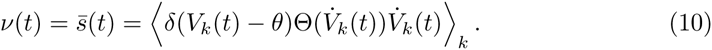

The underlying ensemble of the population is captured by the distribution of voltages and voltage time derivatives at a given time, 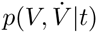. When each neuron’s state is identically and independently distributed, the average over *k* neurons is an average over this distribution at fixed *t*,

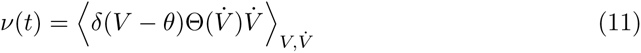

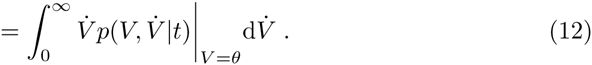

This time-varying expectation value over the statistics of the voltage dynamics in the population is the central time-domain quantity in the response theory for neuronal populations. It is in general analytically intractable.

Subthreshold dynamics can be approximately linear and the many, weak inputs to each neuron can permit a diffusion approximation to a Gaussian process input. In this situation, a model of voltage dynamics that omits the nonlinear voltage reset gives a voltage statistics that is also Gaussian and can be treated analytically. This is the Gauss-Rice approach, to our knowledge first published by Jung [57], where it was used to calculate correlation functions. The first application of the approach to dynamic gain appears in Supplementary Note 3 of ref. [37]. Note that the lack of a voltage reset required for this approach restricts its range of applicability (see Methods and Discussion).

#### Dynamic gain for the mean channel of a population of Gauss-Rice neurons

Here, we derive the dynamic gain using the Gauss-Rice approach, similar to [19, 37, 42]. The emphasis of the derivation here differs in that, first, we go directly to the linear response by linearizing around the mean voltage, and second we express the results in terms of the voltage transfer function to emphasize the additional filtering of the voltage dynamics by the spike threshold. The resulting expression, Eq. (22), applying to a generic population of neurons specified only by the Gaussian statistics and frequency response of their mean voltage dynamics, simply adds a first-order highpass filter to the frequency response of the voltage. In units of the differential correlation time, *τ_s_*, the characteristic time, *τ_c_*, of this high pass depends only on the stationary firing rate, *v*_0_.

Because at zero-lag the voltage and its time derivative are uncorrelated for a stationary variance channel, 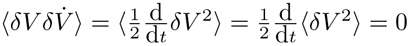, the Gaussian probability density function of the voltage dynamics factorizes over *V* and 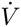,

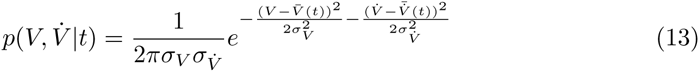

where 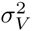 and 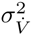 are the respective variances. Substituting this expression into Eq. (12), we obtain

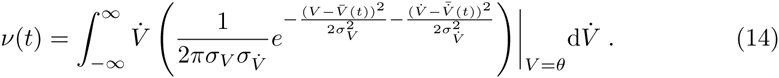

This expression can be computed in terms of error functions to obtain the full nonlinear dynamic response, e.g. for the Gauss-Rice LIF neuron model [19,37,42].

For a transparent analytical treatment of the mean channel in the fluctuation-driven regime we consider the linear response. That is, for case of weak mean input we expand, for each time *t*, this expression in terms of the resulting weak deviations to the ensemble mean voltage 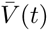 and to its derivative 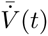. To linear order,

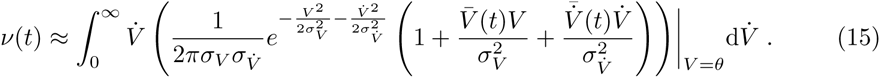

Solving the integral, one obtains the linear response in the mean signal channel,

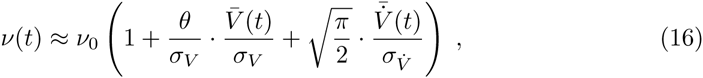

where *v*_0_ is the stationary firing rate attained in the absence of modulation around the mean input current, *I*_0_,

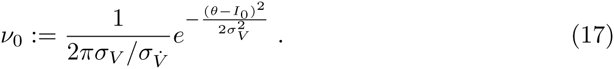

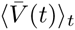 is offset by *I*_0_ and since *I*_0_ ≪ *θ* in the fluctuation-driven regime we set *I*_0_ to 0 without loss of generality, so that 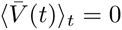 This expression can then be rewritten using only two quantities: the differential correlation time 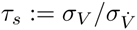 and the size of voltage fluctuations relative to threshold, σ:= σ_V_*/θ*,

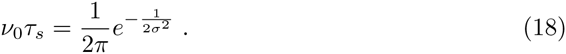

*τ_s_* is the width of the quadratic approximation to the correlation function around zero delay. It serves as the summary timescale determined by the joint effects of all intrinsic timescales and we study in detail this dependence for the GIF model in a later section. *τ_s_* thus provides a natural time unit by which to measure the rate of output spikes, *v*_0_, as a function of the relative voltage fluctuations, σ. *v*_0_*τ_s_* is then interpreted as the number of spikes in a correlated window of voltage trajectory, and according to Eq. (18) rises with σ, saturating for large *σ* at (2π)^−1^ < 1. Fluctuation strength is less than the voltage difference between resting and threshold for most physiological conditions, σ ≲ 1, in which case the useful bound, 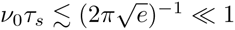, holds. (Large output firing rates can nonetheless be achieved so long as the voltage correlation window, *τ_s_*, is short enough to maintain *v*_0_*τ_s_* ≪ 1.) Spike-generating voltage excursions are thus on average well-separated in time so that the produced spiking exhibits low temporal correlations.

According to Eq. (16), we can then identify *v*_1_(*ω*) as the finite frequency component of its Fourier transform,

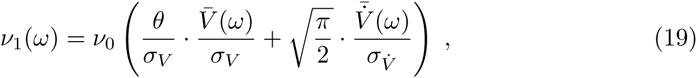

where we note that our definition of *v*_1_(*ω*), Eq. (16), that has the amplitude of the input modulation, *A*, factored out implies that *A* has been factored out of the voltage response. All response quantities are implicitly defined as these *A*-independent versions. This expression can be simplified further by pulling out the time-derivative operator. In the Fourier domain, this is just multiplication by *iω* so that the 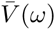 factors out and calculation of *v*_1_ (*ω*) requires only the first two voltage moments, as any statistic derived from a stationary Gaussian process should. 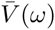 is the mean voltage response and the variances, 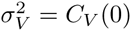 are computed from the correlation function of the stationary unperturbed voltage correlation function, 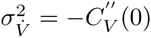 obtained from the voltage noise spectrum *δV*(*ω*). The latter provides only the variances, and so in the space of correlation functions, only directions along which these quantities change affect the rate response [43]. The relative response can then be written

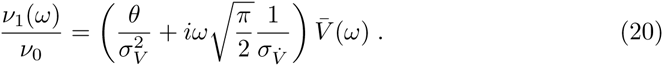

We can re-express it using *τ_s_* and σ,

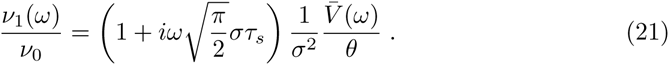

The ensemble response of a population of Gauss-Rice neurons to a small modulation in the mean input is thus simply a first-order high pass filter of the ensemble mean voltage response with characteristic frequency 1/τ_c_, with *τ_c_* defined as

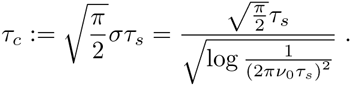

where we have removed σ with 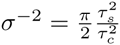, obtained from Eq. (18).

The relative linear rate response is then

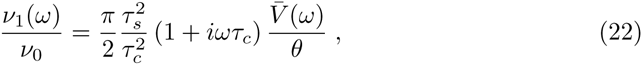

where the dependence on *v*_0_*τ_s_* is concealed in the definition of *τ_c_*. Thus, in units of *τ_s_*, the high pass filter resulting from crossing the spike threshold is proportional to 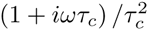 with

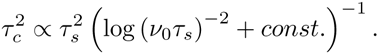

From Eq. (22), we see that the characteristic frequency, 1/*τ*_c_, shifts to lower values for larger output firing rates, as the prefactor, 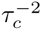, further attenuates the low frequency response. One consequence is that the effect of the low pass voltage characteristics are made negligible by the differentiating action of the spike at high firing rate.

The dynamic gain of this complex-valued linear rate response function is its modulus,

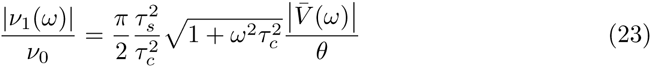

here normalized by the stationary rate, *v*_0_. Taking out a factor of *v*_0_ of Eq. (9), we see that the strength of the linear term and thus the quality of the linear approximation of the response is then controlled by the size of the right hand side of Eq. (23) relative to 1. The effect of this spiking filter contributes a factor that scales as 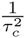 when *τ*_c_ ≪ 1 (for a bounded range of relevant input frequencies) so the linearity assumption is better at larger values of *τ_c_*, which means larger values of *v*_0_*τ*_s_. The quality of the approximation will also depend on the size of 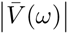. We also note that focusing on the linear response neglects boundedness features of the population firing rate such as its non-negativity. Nevertheless, once a voltage dynamics is specified, Eq. (23) gives the explicit dependence of the dynamic gain on the underlying parameters of the single neuron model.

### Derivation of the dynamic gain of a population of Gauss-Rice GIFs

In this section, we take the general result of the previous section, Eq. (23), and go through its explicit calculation for a population of Gauss-Rice GIF neurons to obtain the result Eq. (41). A work taking a similar approach, partly inspired by this work, though with with less intermediate analysis has recently appeared [25]. Our novel findings arise from an exhaustive characterization of the parameter dependence across the phase diagram of the voltage response, Fig. 3. We calculate the current-to-voltage filter, expressing it in each of the three representations listed in Table 1, Eqs. (25,26,27) respectively. We show (Fig. 4) how the the low pass component of the filter undergoes a qualitative change from second-order low pass to first-order low pass to resonant as *Q_L_* is increased. We find the voltage resonance condition, 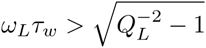 where the resonance has a contribution from slow adaptation and from the frequency, Ω. Either can exist without the other (Fig. 5). We then compute the voltage correlation function, Eq. (37), whose envelope depends on the relaxation time, *τ_r_* (Fig. 6). From this, the variances are calculated and an expression for the differential correlation time, *τ_s_*, Eq. (39) is obtained. We show a characteristic dependence on the ratio *τ_w_/τ_I_* (Fig. 7). Finally, we show in Fig. 8 how the stationary firing rate has unimodal dependence on the time constants, *τ*_v_ and *τ_I_*, monotonic rise with input variance, 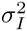, and monotonic decay with intrinsic frequency, Ω.

**Figure 3.**
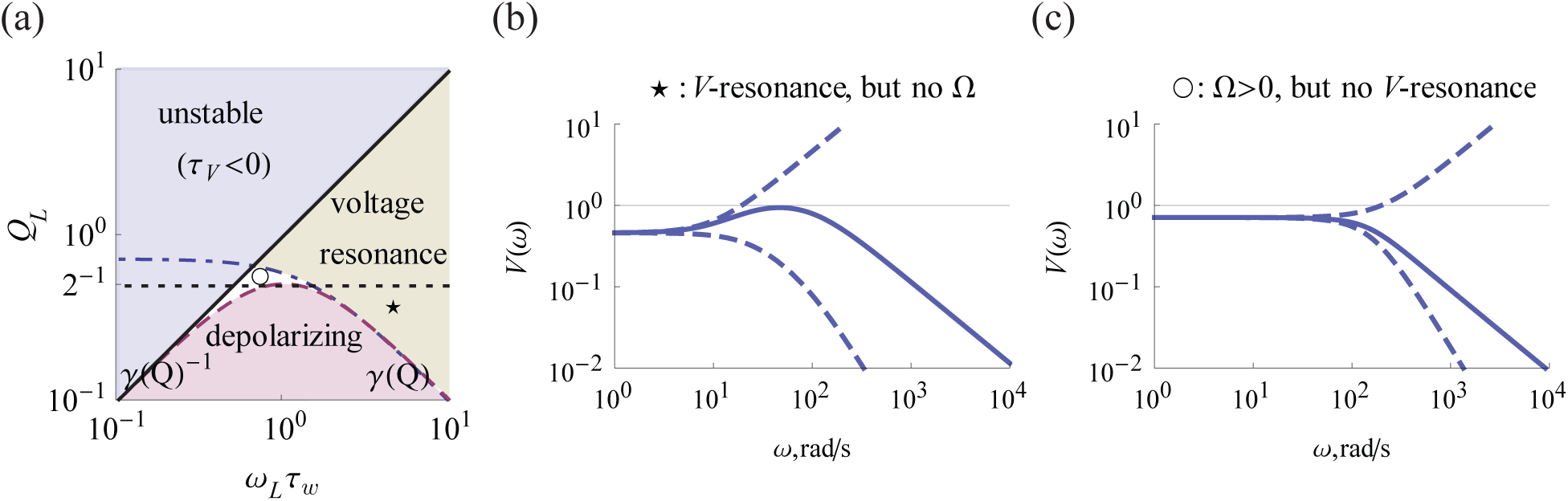
Regimes of the current-to-voltage transfer function. (a) Phase diagram of the transfer function. The region of depolarizing *w* (low frequency amplifying, *V_low_* > 1) is shown in purple and voltage resonance in green. The filter is unstable in the blue region. An intrinsic frequency exists, above the dotted line, *Q_L_* = 1/2. Note that there is a region with *Q_L_* > 1/2 and no voltage resonance, and vice versa. The star and circle denote the example values of (*w_L_*τ*_w_*, *Q_L_*) used in (b) and (c), respectively. (b) An example of the current-to-voltage filter in the case of resonance with no intrinsic frequency (*τ_V_* = 10, *τ_w_* = 100, *g* = 1.2). (c) An example of the current-to-voltage filter in the case of no voltage resonance despite the existence of an intrinsic frequency (*τ_V_* = 10, *τ_w_* = 5, *g* = 0.5). The rising and falling dashed lines in (b) and (c) denote the contributions of the high pass, 1 + *iwτ_w_*, and the low pass, 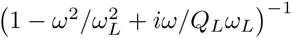, respectively. Their combination forms the current-to-voltage filter, which are shown as solid lines.

**Figure 4.**
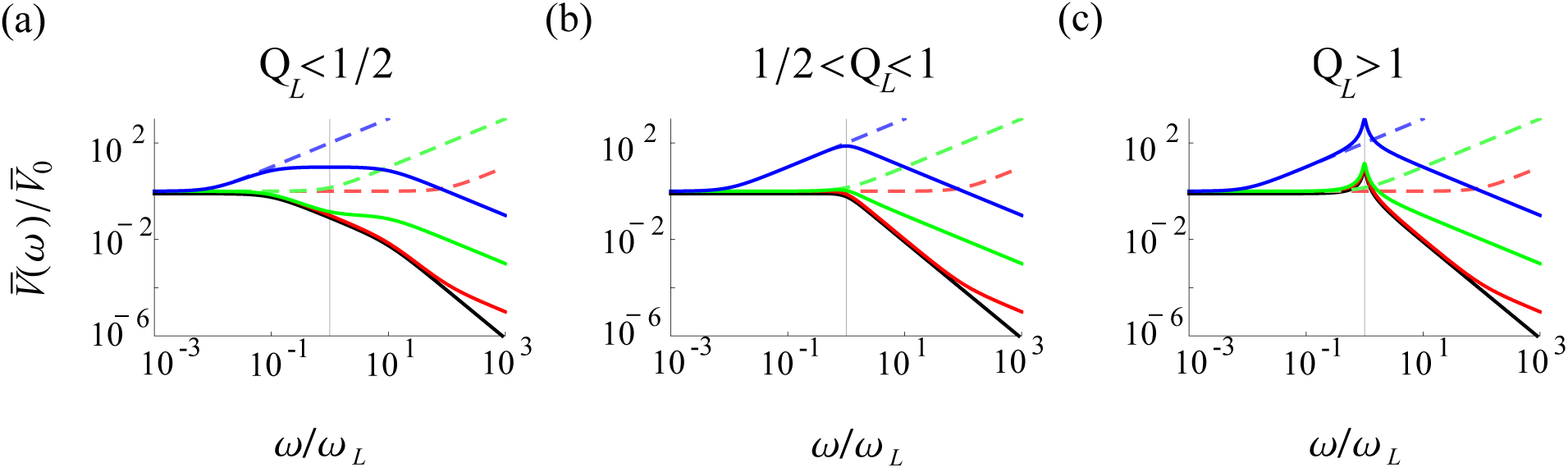
The qualitative shape of voltage response depends on *Q_L_*. Here we classify the current-to-voltage filter shapes shown as colored solid lines in (a), (b), and (c), which show the three *Q_L_*-regimes with respective examples for *Q_L_* = 0.1,0.75,10. In each plot, the high pass component of the voltage response is shown as the colored dashed lines, one for each of three representative values of its characteristic frequency, *w_L_τ_w_* = 10^−2^ > γ(blue), *w_L_τ_w_* = l(green), and *w_L_τ_w_* = 10^−2^ < γ^−1^(red). The solid black line is the low pass component of the voltage response. For the regime shown in (a), the green case can not be achieved when *w* is hyperpolarizing (*g* > 0) and the example red case cannot be achieved because it violates the stability condition *Q_L_ < w_L_τ_w_*.

**Figure 5.**
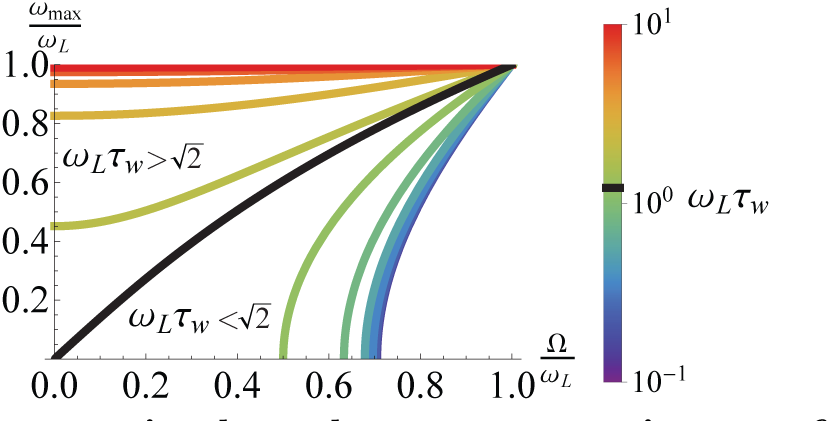
A resonance frequency emerges in the voltage response in one of two ways depending on the intrinsic timescale. For slow intrinsic current 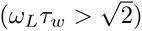, a response exhibiting a maximum at *w_max_* already exists at Ω = 0. For fast intrinsic current 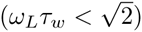, a resonance emerges at finite Ω, whose value converges for vanishing *w_L_τ_w_*.

**Figure 6.**
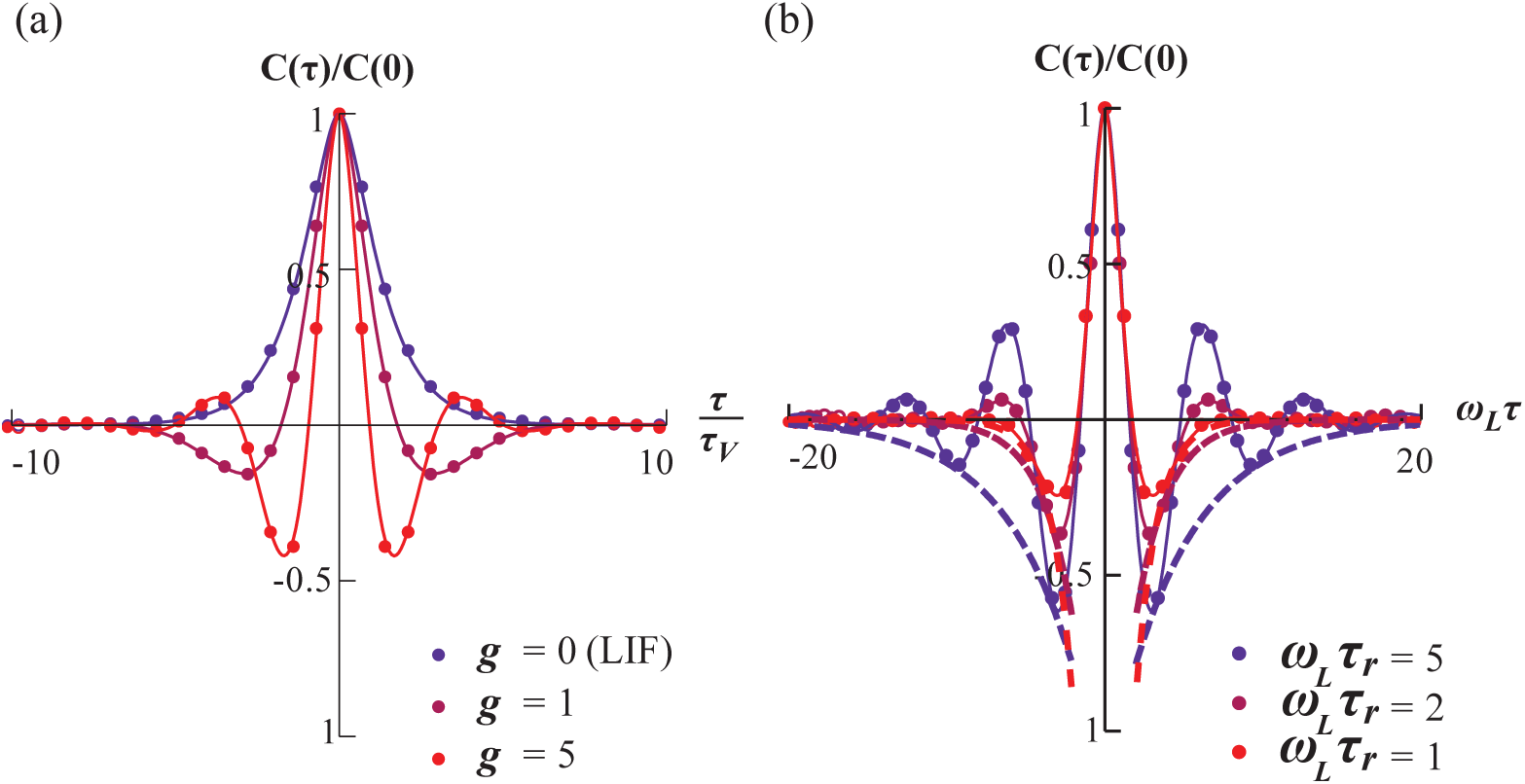
Emergence of oscillatory behavior in the voltage dynamics and a consequent ringing appears in the voltage correlation functions. (a) The frequency of the ringing increases with the strength of the intrinsic current (*g* = 0,1, 5 shown; *τ* in units of *τ_v_*). (b) The envelope of the ringing widens with *τ_r_* (dashed lines are 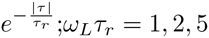 shown and *τ* in units of 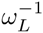). Lines are Eq. (37); dots are numerics.

**Figure 7.**
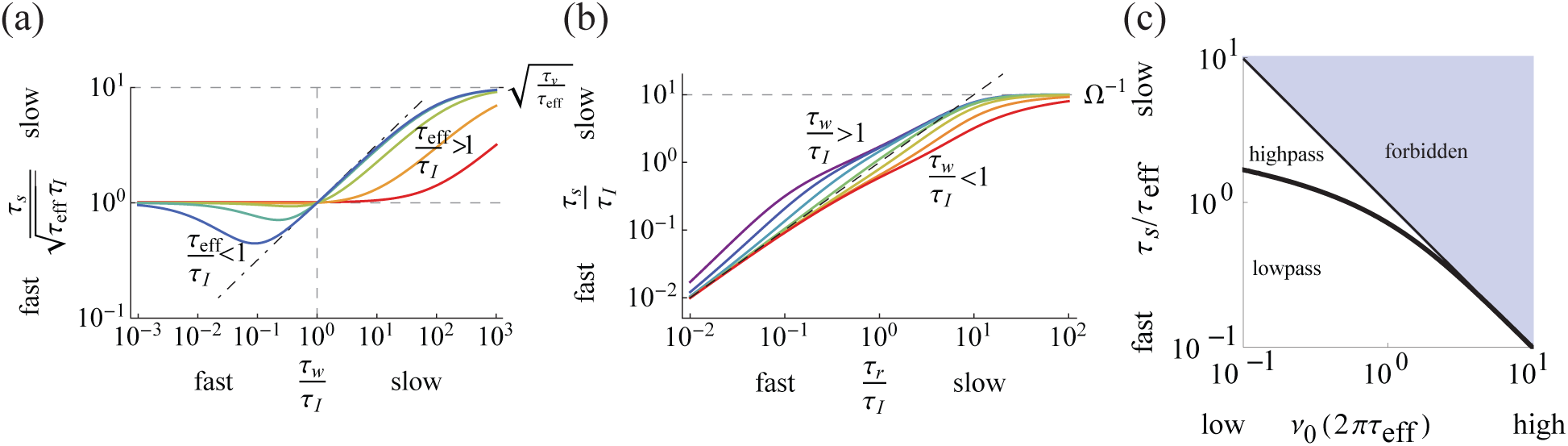
Differential correlation time depends on intrinsic parameters. (a) *τ_s_* increases (not always monotonically) with *τ_w_*. For the sake of comparison, we show *τ_s_* normalized by its small-*τ_w_* limiting value, 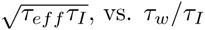 vs. *τ_w_/τ_I_* across *τ_V_/τ_I_* = 10^o^, 10^1^,10^2^,10^3^10^4^ (from blue to red) with *g* adjusted so their large-*τ_w_* limiting value, τ*_V_*/τeff = 1 + g = 10^2^. Shapes are sigmoidal for *τ_eff_*/*τ_I_* > 1 (e.g. green to red) and include an initial dip for *τ_eff_*/*τ_I_* < 1 (blue to green). The dot-dashed line denotes 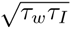. (b) *τ_s_* follows the relaxation time, *τ_r_*, (the dashed line is *τ_s_ = τ_r_*) and saturates at Ω^−1^. Colors indicate the value of *τ_w_/τ_I_* on a logarithmic scale from 10^−1^(red) to 10^1^ (purple). (c) General shape of *τ_s_* vs. *v*_0_. Values in the blue region are forbidden due the maximum rate achievable in a Gauss neuron. The thick black line denotes the boundary between high and low pass.

**Figure 8.**
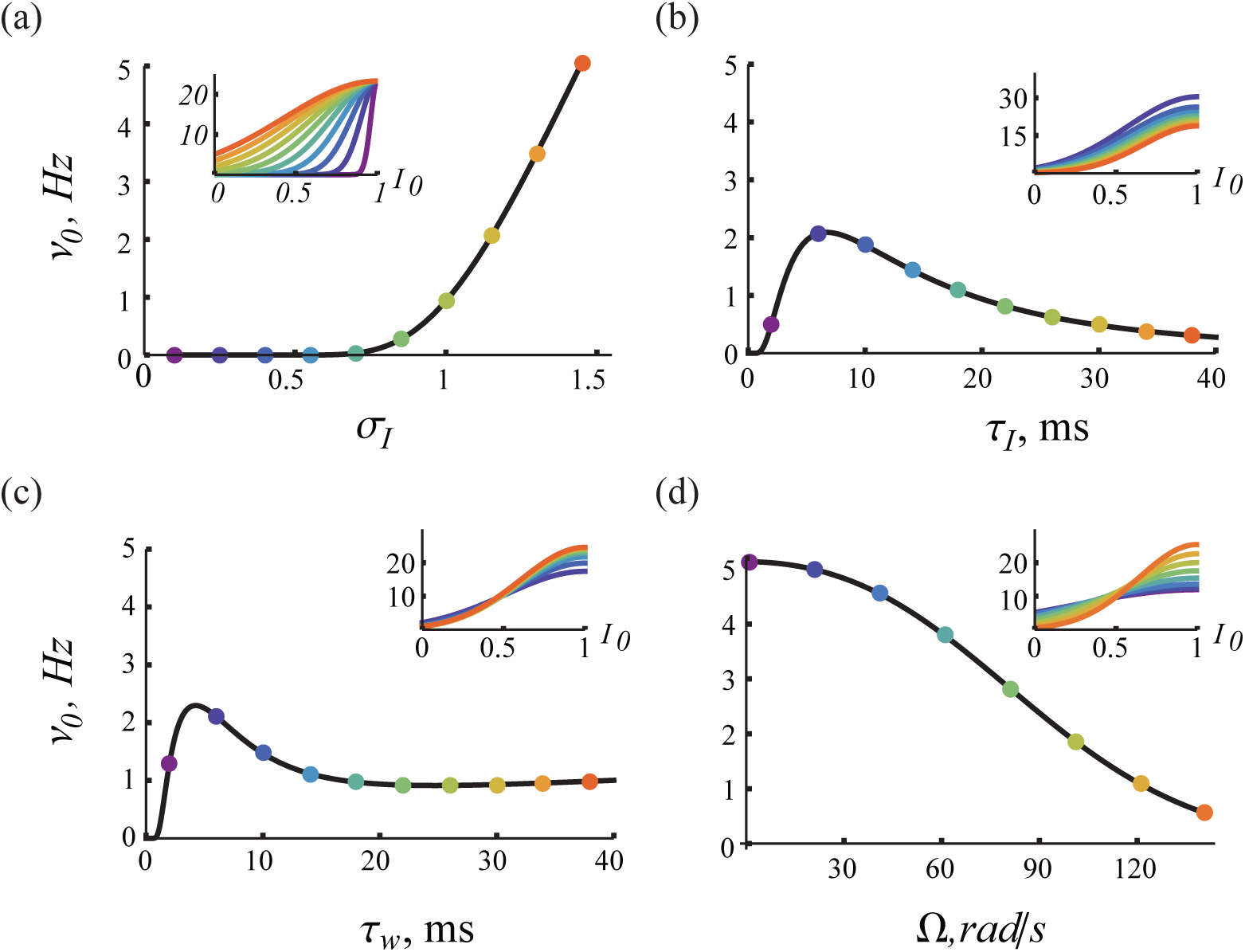
Effect of model parameters on the fluctuation-driven stationary response. The stationary firing rate, Eq. (18) for *I*_0_ ~ 0 (a) increases monotonically with the strength of input fluctuations and (d) decreases monotonically with the intrinsic frequency. Across each of *τ_I_* and *τ_w_* ((b) and (c) respectively), the rate exhibits a maximum. Insets are the mean input dependent expression for the stationary response, Eq. (17), valid in the regime *I*_0_ ≪ 1. Inset color refers to the value of the parameter (*σ_I_, τ_I_, τ_w_* and Ω) at the location of the colored dots in the main plots. Parameters were otherwise set to their default values.

#### Voltage solution

For arbitrary input, *I*(t), the system in the Fourier domain is

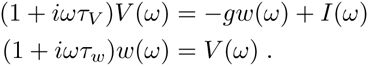

Multiplying the first equation by (1 + *iwτ_w_*) and eliminating *w*(*ω*) one obtains

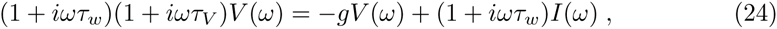

so that the solution for any *g* > −1 is, with respect to the representation of the model by its explicit parameters (*g, τ_v_, τ_w_*),

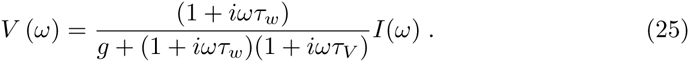

When the neuron exhibits an intrinsic frequency, Ω, we can use 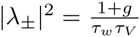 and the definition of the complex eigenvalues by their real and imaginary parts, 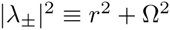 (see Methods), to substitute Ω into the denominator of Eq. (25) after expanding:

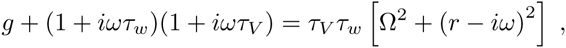

with 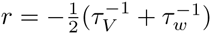 and 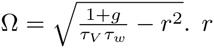 defines the relaxation time of the dynamics, *τ_r_* = −r^−1^. Thus, in the representation of the model based on the implicit time scales, (Ω, *τ*_r_, *τ*_w_), the solution is expressed as

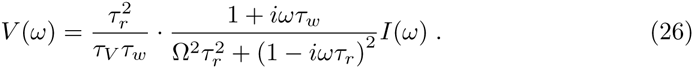

A third convenient representation consists of effective parameters, (*ω*_L_, *Q_L_*, *τ*_w_), determining the shape of the filter

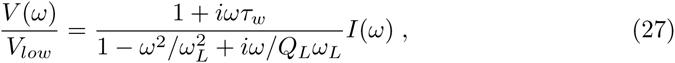

where the second order low pass filter has been re-expressed using its center frequency,

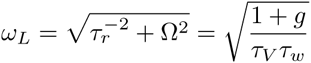

at which its contribution to the gain is its quality factor,

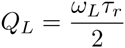

(with 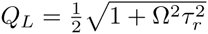 when 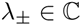), and we have pulled out the broadband voltage response, *V_low_*, attained in the limit *ω* → 0, which gives

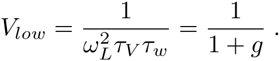

The stability constraint, *g* > −1 is naturally satisfied by *ω_L_* > 0 and keeps *V_low_* finite. With dependence on *τ_v_* removed in the shape representation, we must explicitly add the stability constraint, *τ_V_* > 0, which is expressed using the definition of *τ_v_* in this representation,

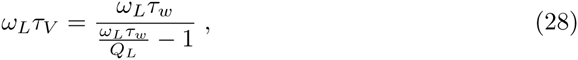

so that the stable regime corresponds to *Q_L_* < *ω*_L_*τ*_w_. *V_low_* is expressed in this shape representation as

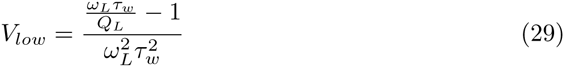

so that V*_low_* > 0 is satisfied by the stability constraint.

Each of the three expressions for the voltage response filter, Eqs (25,26,27), is instructive in understanding the dependence on the contained parameters. To motivate this analysis in the context of population response, we first specify the input, *I*(*ω*). An input oscillation of frequency *ω*_0_ will produce an oscillation in the mean input expressed as 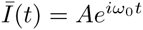. In the frequency domain, the spectrum of the mean input, Ī(*t*), and power spectral density of the noise, *Ī*(*t*), is, respectively,

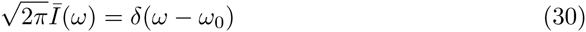

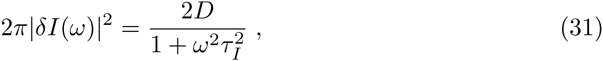

with noise strength, 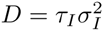, in the latter. Because of the linearity of the dynamics, we can solve the system for mean and fluctuating input separately. In the next paragraph, we employ Eq. (30) to obtain the mean voltage response, and in the following paragraph we employ Eq. (31) to obtain the voltage correlation function.

#### Mean voltage response function

The population mean voltage response, 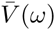, required for Eq. (20) is obtained by inserting the expression for the mean input, Eq. (30), into the voltage solution. For the remainder of the paper, we omit the factor *δ*(*ω* − *ω*_0_) and denote the frequency of the mean input by *ω*. The mean response in the three representations is then Eqs. (25,26,27), respectively, with the *I*(*ω*) factor dropped and with an additional a factor of 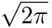

The mean voltage response is given in the filter shape representation, (*τ_V_*, *τ_w_*, *g*), by

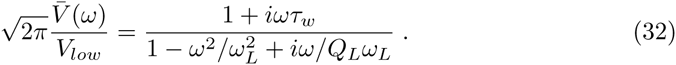

We now go through the analysis of this response using this representation. Using the gain,

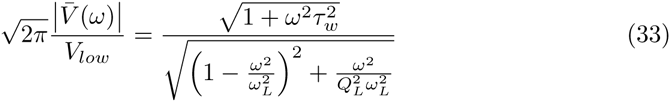

we constructed a diagram of its qualitative features in *Q_L_* vs. *ω*_L_*τ*_w_ (see Fig. 3).

The model exhibits low frequency voltage gain amplification (*V_low_* > 1) or attenuation (*V_low_* < 1) depending on whether *w* is depolarizing (*g* < 0) or hyperpolarizing (*g* > 0), respectively. *ω*_L_ = 0 at *g* = −1 and grows with *g* as 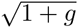. *Q_L_*= *ω_L_τ_r_*/2 also grows with *g*, generating three parameter regions of qualitatively distinct low pass filter gain shapes: *ω_L_τ_r_* < 1,1 < *ω_L_τ_r_* < 2 and *ω_L_τ_r_* > 2. Indeed, in units of *ω_L_*, the shape of the current-to-voltage filter depends only on *τ_r_* and *τ_w_*, and so in the next paragraphs and with reference to Fig. 4, we describe this 2D parameter space completely by considering qualitative differences in the full filter shape across *ω_L_τ_w_* in each of three distinct regimes of *ω_L_τ_r_*. Note that relatively slow and fast intrinsic dynamics is obtained when *Q_L_* is less than or greater than 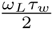, respectively.

For *ω_L_τ_r_* < 1 (see Fig. 4(a)), the low-pass gain contribution can be factored into a contribution arising from two first order low pass filters,

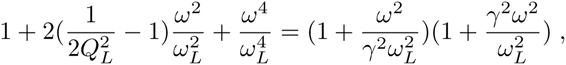

where *γ* = *γ*(*Q_L_*) ≥ 1 is the solution to *Q_L_* = *γ*/ (*γ*^2^ + l). The low pass gain thus begins falling as *ω^−^*^2^ after *ω_L_*/γ and then as w^−4^ after *γω_L_*. The intermediate region, *ω*/*ω*_L_ ∈ (*γ*^−1^, *γ*), is given by the inequality *Q_L_* <γ/ (*γ*^2^ + l) and disappears as *Q_L_* approaches 1/2 where *γ* and *ω_L_τ_r_* approach 1. The region of depolarizing *w* (*g* < 0) shown in Fig. 3 satisfies 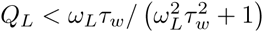 in this representation, whose solution in *ω_L_τ_w_* is also the range (*γ*^−1^,γ). Thus, response shapes in this intermediate region (see Fig. 4(b)) are only achievable by depolarizing *w*, and *w* must be depolarizing for any response exhibiting such shapes. Consequently, the three qualitatively distinct shapes of the current-to-voltage filter for *ω_L_τ_r_* < 1 are determined by the location of *ω_L_τ_w_* relative to 1/*γ* and γ, with the middle regime, (γ^−1^,γ), only achievable for depolarizing *w*. For *ω_L_τ_w_* > γ, the filter first rises with *ω* after 1/*τ_w_*, is flattened at *ω_L_*/γ, and then falls after γ*ω_L_*. The result is an intermediate, raised plateau of width (γ − γ^−1^) *ω_L_*. The condition for this voltage resonance is 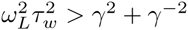 or in terms of *Q_L_*, *Q_L_* > (2 + *ω_L_τ_w_*)^−1/2^. For 1/γ < *ω_L_τ_w_* < γ, the response attenuates first and so the plateau is now an intermediate, downward step of width (γ − 1)*ω_L_*. For *ω_L_τ_w_* < 1/γ, there is only low pass behavior and the high pass only acts to pull up the *ω*^−4^-falloff up to a *ω*^−2^-falloff. As *ω_L_τ_r_* approaches 1 from below, γ also approaches 1, and the qualitatively distinct region between *ω_L_/γ* and γ*ω_L_* shrinks as the two roots coalesce into one and the low pass expression forms a perfect square. In the case that *ω_L_τ_w_* > γ, this leave a well-defined maximum located just before *ω_L_*. The slight offset arises simply because the second order low pass begins falling significantly before *ω_L_* at *ω_L_τ_r_* = 1.

For 1 < *ω_L_τ_r_* < 2 (see Fig. 4(b)), the impact of the high-pass on the shape of the filter is determined simply by whether its characteristic frequency is above or below *ω_L_*. For *ω_L_τ_w_* > 1, the plateau existing for *ω_L_*τ*_r_* < 1 becomes a flat-topped peak in the gain with a maximum again slightly lower than *ω_L_*. Otherwise, the behavior is low pass. Note that *ω_L_τ_r_* > 1 is also where the intrinsic frequency exists. However, this property does not contribute to a resonance until *ω_L_τ_r_* > 2. Indeed, the resonance here, as in the regime *ω_L_τ_r_* < 1, arises solely from a high pass attenuation of low frequencies sculpting a peak from a low pass, and comes alongside a region, *Q_L_* < (2 + *ω_L_τ_w_*)^−1/2^, that lacks resonance. This latter region is upperbounded in general by 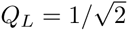, and specifically for stable filters by 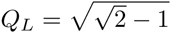 so that above these values of *Q_L_* all filters are voltage resonant.

For *ω_L_τ_r_* > 2 (see Fig. 4(c)), by definition a resonant peak emerges in the low pass filter. If *ω_L_τ_w_* < 1, this contributes a *de novo* resonance in the current-to-voltage filter located near *ω_L_*. Otherwise, it simply acts to sharpen the existing resonance that appears progressively over 0 < *ω_L_τ_r_* < 1, and again with a peak slightly to the left of *ω_L_*.

Of the two mechanisms for resonance just described, the contribution of first ‘sculpting’ mechanism leads to a linear increase in the response height and input frequency range of elevated response with *τ_w_*, i.e. with the slowness of the intrinsic dynamics, for the reason that the low frequency amplification continues over a broader range the further *ω_L_*/γ and 1/*τ_w_* are apart. This amplification in the relative response is actually over-compensated by a broadband attenuation with *τ_w_*, so that the actual effect is the carving out of a resonant peak using adaptation, i.e. a low frequency attenuation of an otherwise low pass filter.

The second low-pass resonance mechanism emerges in the expression when the low pass filter exhibits a maximum, which itself emerges when the two low pass characteristic times of the low pass coalesce. From the point of the view of the voltage dynamics, this occurs from a sufficiently strong and negative feedback interaction between *v* and *w*, whose timescales are sufficiently similar so that the delayed feedback is constructive. In the time domain voltage solution, this occurs when the two eigenvectors align. The height of the resonant response grows linearly with *τ_r_* (with range of elevated response fixed) because there is less dissipation.

These two resonance mechanisms contribute to the height of the response at *ω_L_*,

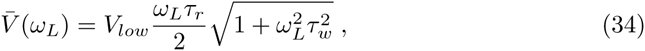

which is resonant by definition if it is greater than *V_low_*. The condition for voltage resonance is thus 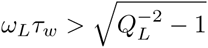 and the relative ratio of their contributions is 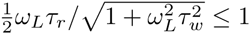 so that at a given *ω_L_τ_w_* the sculpting mechanism always contributes more gain than the intrinsic frequency mechanism. Indeed, this sculpting can exist in the absence of an intrinsic frequency (*ω_L_τ_r_* < 1), so long as the intrinsic dynamics is slow enough. Conversely, even with an intrinsic frequency (*ω_L_τ_r_* > 1), the response can lack a resonance if in addition *ω_L_τ_w_* < 2, demonstrating that an intrinsic frequency is not a sufficient condition for resonance. These two cases become apparent in a plot of the resonance frequency as a function of the intrinsic frequency (Fig. 5), where we observe x- and y-intercepts because of the preexisting or absent resonance, respectively. The location of the maximum converges to 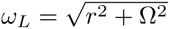 for *ω_L_τ_w_* ≫ 1, which itself converges to Ω for Ω*τ_r_* ≫ 1. For smaller values of *ω_L_τ_w_*, the location converges to a value slightly larger than *ω_L_*.

We will make use of the representation of the current-to-voltage filter in terms of (*ω_L_, τ_w_, Q_L_*) to understand the full response. What is left to calculate, however, is the voltage correlation function.

#### Voltage correlation function and the variances, 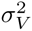 and 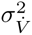

For the correlation function of *V*, we perform the calculation in the implicit representation and add Eq. (31) to the modulus squared of Eq. (26),

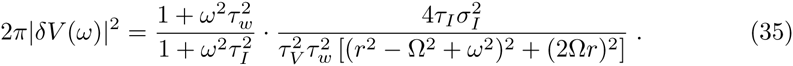

The auto-correlation thus requires computing an inverse Fourier transform integral of the form

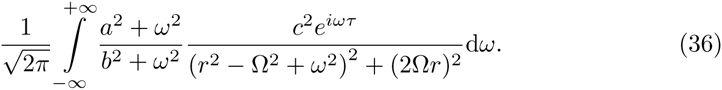

The result is

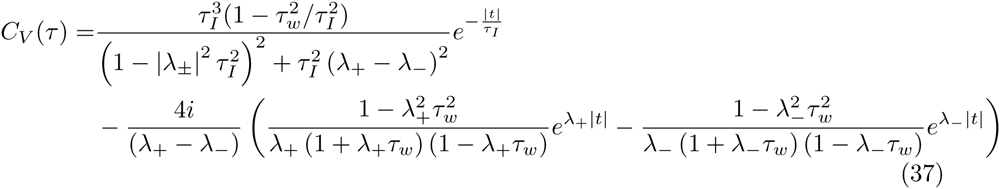

where λ_±_ are the eigenvalues of the voltage dynamics, Eq. 57, and the units are [Time^3^]. The correlation has two components, one decaying with *τ_I_* and the other with *τ_r_*. The first component is strongly suppressed for *τ_r_*/*τ_I_* ≪ 1. The second component exhibits damped oscillations within the exponential envelope with frequency Ω. Examples are shown in Fig. 6. for increasing *g* and *τ*_r_. Note that variation in *g* affects the width of the function around 0-delay while it is fixed over a variation in *τ_r_*. These results were checked against numerical autocorrelation functions computed from the voltage time series output of the numerically implemented model. The correspondence is excellent.

In the model representation, the variance of the voltage and that of the time derivative of the voltage are given by

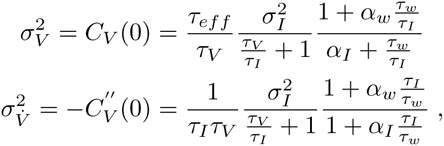

where, for notational convenience, we have defined

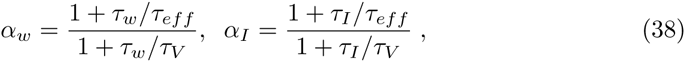

and *τ_eff_* = *τ_V_*/(1 + *g*) (Eq. (2)) is the maximum speed over *τ_w_* of the voltage kinetics, approached when *τ_w_ ≪ τ_V_* by the tonic conductance change induced by *w*. For the LIF (*g* = 0), *τ_eff_ = τ_V_, α_I_ = α_w_* = 1 and the variances simplify to

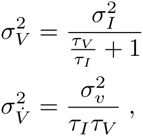

from which the differential correlation time *τ*_s_ for the LIF can be read off as 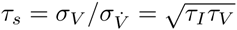. We consider this quantity more generally in the next paragraph.

In the intrinsic representation, the variances can be written as

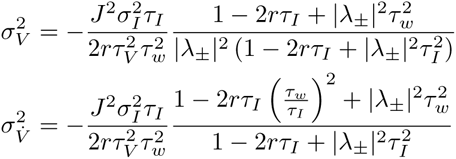

where |λ_±_|^2^ = *r*^2^ + Ω^2^, and *r* < 0 ensures that the values are positive. Note that the only difference between the expression for the two variances is a factor of 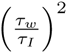 and a factor of 1/|λ_±_|^2^. In all representations, the influence of intrinsic kinetics set by *τ_w_* is negligible when *τ_w_* is near the input timescale, *τ_I_*. Even then, *w* affects the variances via *g* or Ω.

#### The differential correlation time and the stationary response

From the correlation function providing the variances, the differential correlation time is calculated with 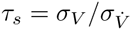. The Gauss-Rice GIF differential correlation time for the model representation is

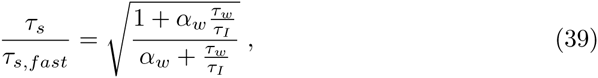

where the limiting value of *τ_s_* for *τ_w_* smaller (larger) than all other timescales is, respectively

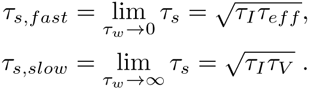

The ratio of slow and fast limiting values is 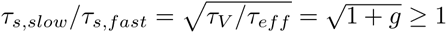, so that *τ_s_* increases over the full range of *τ_w_/τ_I_*. In particular, the curves of Eq. (39) have a characteristic shape for the non-trivial (*g* ≠ 0) cases. We focus on the hyperpolarizing case. In the left panels of Fig. 7, we plot some example shapes of *τ_s_/τ_I_* vs. *τ_w_/τ_I_* over a range of *τ_eff_* < *τ_V_*. Referring to that figure, for *τ_eff_* < *τ_I_*, the curves monotonically interpolate between the limiting values, with the abscissa value at half-maximum increasing linearly with *τ_V_/τ_I_*. With *τ_w_/τ_I_* increasing from 0, *τ_s_* first drops from 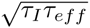 to a minimum (whose depth grows with *g*) and then rises into a 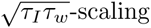 regime around 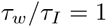, where it passes through the same value as that attained in the limit*τ_s,fast_* and then eventually saturates for *τ_w_*/*τ_I_* ≫ 1 at its maximum, 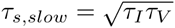. Thus, for *τ_w_/τ_I_* → ∞ to the *g* ≠ 0 case is equivalent to the *g* = 0 case and we conclude that any novel features attributable to the extra degree of freedom are washed out in this limit by the relatively slow intrinsic dynamics. As discussed in the Methods, the validity of the no-reset approximation lies around *τ_s_/τ_j_* ~ 1, implying that *τ_v_* ≳ *τ_I_*. When *τ_eff_* ~ *τ_I_*, the approximation is valid across *τ_w_ < τ_I_* and for other *τ_eff_* < *τ_I_* only in ranges around the value of *τ_w_ > τ_I_* for which *τ_s_ ~ τ_I_*. We also find for relatively slow intrinsic dynamics that *τ_r_ ≲ τ_s_*, for τ_s_ ≤ Ω^−1^. When Ω exists, we can write *τ_s_* as

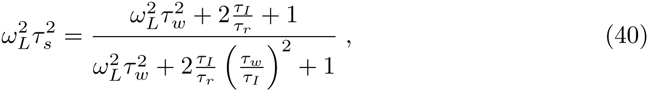

where 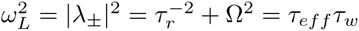, is the center frequency analyzed in the previous section. In the implicit representation and as a function of *τ_r_* (see Fig. 7), *τ_s_* grows faster and slower than linear for *τ_w_/τ_I_* less than or greater than 1, respectively, and passes through 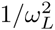 when *τ_w_/τ_I_* ~ 1, finally saturating at Ω^−1^. Up to this saturation level, *τ_s_ > τ_r_* for *τ_w_ < τ_I_*, so that the condition *v*_0_*τ_s_* ≪ 1 implies that *v_0_τ*_r_ ≪ 1 and the approximation to reset dynamics is valid. In the case *τ_w_ > τ_I_*, the range of *τ_r_* over which the *v_o_τ_r_* ≪ 1 validity constraint is not already covered by the *v*_0_*τ_s_* ≪ 1 built-in constraint is centered around *τ_w_* = Q^−1^ and grows in size with *τ_w_/τ_I_*.

Next, we compute the stationary firing rate of the neuron model Eq. (1) as a function of the two input parameters and the two intrinsic parameters. It is shown in Fig. 8. We focus on the parameter dependence at *I*_0_ = 0. The model’s stationary response to increased input noise exhibits a cross-over from silence to linear growth around σ*_I_* ~ *θ*, simply due to the higher propensity of threshold crossings. In subsequent analyses in this paper, we explore the parameter dependence at fixed stationary output firing rate by adjusting the input variance accordingly (see Methods for this mapping). The rate dependence at *I*_0_ is similar for both *τ_I_* and *τ_w_*, growing from zero at vanishing time constants to a maximum located just below the membrane time constant. While the rate decays with increasing *τ_I_*, it seems to saturate and even rise for slowly with *τ_w_* for *τ_w_* > *τ_V_*. The stronger the flow of the dynamics around the resting state at *I*_0_, the more the voltage fluctuations are dampened so that the the firing rate decreases with Ω. As for the *I*_0_-dependence, we see that all curves rise monotonically simply because the average voltage moves closer to the threshold.

#### Expression for the complex response function

With the variances and the mean voltage response in hand, we can write down the complex linear frequency response,

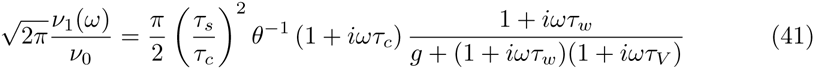

This biquad filter is composed of two-step cascade of a combined 1st-order high-pass and 2nd-order low-pass current-to-voltage filter followed by a first-order high-pass voltage-to-population firing rate filter. In the remaining part of the paper, we analyze the properties of this filter.

### Analysis of the dynamic gain function of a GIF ensemble

In this section, we characterize the qualitative features of the response function, Eq. (41), again with a focus on completeness. We first show that the high and low input frequency limits of the response constrain the parameter sets that can achieve high and low pass behavior and we give an expression, Eq. (45) of the critical stationary rate separating these two regions in terms of the other parameters. We then reparametrize the expression for the response, Eq. (47), using the height of the response at its center frequency, 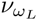 and high frequency limit, *v_∞_*, both relative to its low frequency limit. The two-dimensional shape parameter space give responses with a peak, dip or step at *ω_L_* whose width varies with *Q_L_*. The additional high or low pass nature of the filter give six classes of filter shape. The constraint of stable voltage dynamics restricts the area accessible to the model 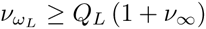

### The *ω* → 0 and *ω* → ∞ to limits simply determine a high/low pass criterion

The matched order between the high and low pass filter components of Eq. (41) implies that there are finite limiting values of the dynamic gain at low and high input frequencies,

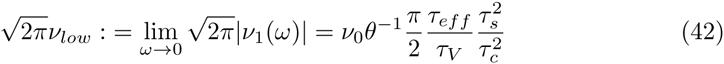

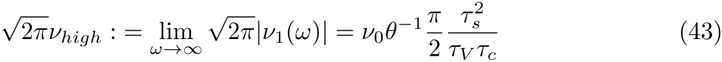

with the size of *v_hioh_* relative to *v_low_*, 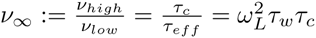. We note that both *v_low_* and *v_high_* can be written without explicit dependence on the intrinsic timescale, it influences the limiting values only by setting the value of *τ_s_* in the way demonstrated in the previous section.

*v_low_* scales the above filter shapes up or down and itself scales down linearly with *τ_eff_*/*τ_V_* and thus with *g*. The boundary in the parameter space between low and high pass is defined implicitly by *v_∞_* = 1 providing the simple criterion for low or high pass behavior as whether *τ_c_* is below or above *τ_eff_* respectively. The high pass behaviour for large *g* or *Q_L_* is not due to an increase in *v_high_* (in which *g* does not appear) but in fact a consequence of the low frequency attenuation. Recalling that the approximation to a hard threshold keeps the response flat to arbitrarily high frequencies, while in fact it eventually decays (beyond *f_limit_*, as discussed in the Methods section), the high pass case here implies a large elevated high frequency band up to this cut-off, while the low pass condition implies a large intermediate downward step. The low/high pass criterion implies a critical relative variance 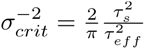, and in turn the critical output firing rate,

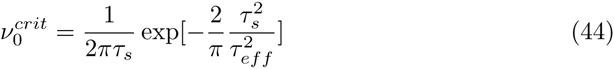

at which the response changes from low to high pass. Both of these values are intrinsic properties of this model whose dependence on the input relies only on the units of time taken. For 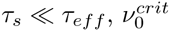 diverges as 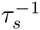 For *τ_s_* ≫ *τ_eff_*, it falls off as 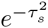 In Fig. 7(c), we plot *τ_s_* as a function of *v_0_*. One can now use the plot in this figure to determine the high or low pass behavior for a given *τ_w_*/*τ_I_* and *v_0_*. For example, when *τ_s_* < τ*_eff_* (attained for instance with small *τ_w_* and large *τ_V_*/*τ_eff_*), there is only low-pass behavior due to the divergence of 1/2*πτ*_s_. The high pass region nevertheless grows quickly with *τ_s_* > *τ_eff_*.

The low and high input frequency limits become independent of *τ_s_* when time is expressed in those units. Nevertheless, we can still write the critical condition independent of *τ_s_* when expressing time in units of *τ_I_* by combining Eqs. (42) and (43) and eliminating *τ_s_* altogether by substituting in the expression for 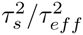 (*cf.* Eq. (44)) to get the high-low pass condition explicitly and solely in terms of the four timescales: *τ_w_*, *τ_eff_*, *τ_V_*, and 1/*v*_0_ (the latter value chosen by setting *σ_I_* appropriately using Eq. (60)). Setting any three of these determines the critical value of the remaining one above, across which the model changes from high to low pass behavior. For example, when time is measured in units of *τ_I_*, we have

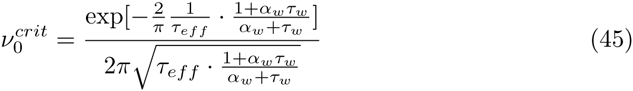

For *g* = 0, we have *α_w_* = 1 and *τ_eff_* = *τ_V_* and this reduces simply to 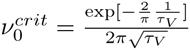, the expression presumably underlying results for the LIF in [42].

As for the limiting behavior of the phase response, Φ(*ω*),the model gives zero delay for both high and low frequencies. At low frequencies, this is because the input changes slowly so that the model dynamics can directly follow the oscillation. At high frequencies, the return of the lag to 0, just like the flat high-frequency gain, is an artifact associated to the hard threshold.

#### There are six qualitatively distinct filter shapes

When *g* = 0 (LIF), the filter, Eq. (41), simply reduces to single order. The intermediate behavior is then only the respective monotonic decay or rise beginning and ending around the smaller and larger of the two characteristic frequencies.

For *g* ≠ 0, the voltage modulation by the current, *w*, comes into play. To analyze the effect of the high pass voltage-to-spiking filter on the current-to-voltage filter we employ a similar exhaustive characterization as was done above in the analysis of the current-to-voltage filter, i.e. by going through all the cases arising from distinct orderings of the characteristic times of the components of the combined filter. The ordering can give simple information about the filter shape. For instance, any contribution of the voltage-to-spiking filter to the qualitative behavior of the complete filter beyond just low or high pass requires that 1/*τ_c_* be no larger than either *ω_L_* or 1/*τ_w_*. Otherwise, the only effect of the spiking is to flatten the high frequency response beyond 1/*τ_c_*. In general, however, there are many possible shapes. To further facilitate the classification of these shapes, we present a single parameter space representation in which they are all simply mapped.

For this general case, we can introduce the relative quality factor for the full filter, 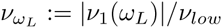. The response then depends on the five shape features, *v_low_, v_high_, w_L_*, *Q_L_*, and 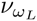 Denoting ξ = *τ_w_*/*τ_c_*, so that 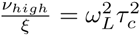 and 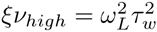, we can re-express the response function as

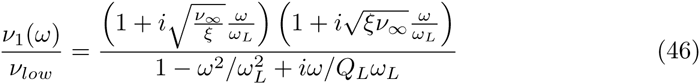

with dynamic gain

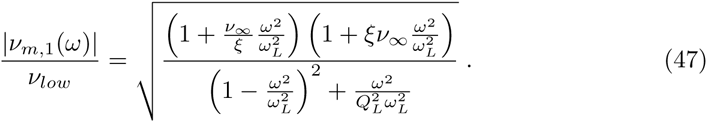

When 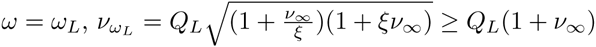, which implicitly defines *ξ* in terms of 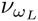 *Q_L_* and *v_high_* and closes the representation. Indeed, with time in units of 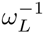 and gain values relative to *v_low_*, the shape of the filter depends only on this triplet: each of the six regions in 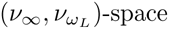 defined by the boundaries *v_∞_* = 1, 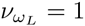, and 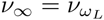 provides filters of a qualitatively similar class (see Fig. 9).

**Figure 9.**
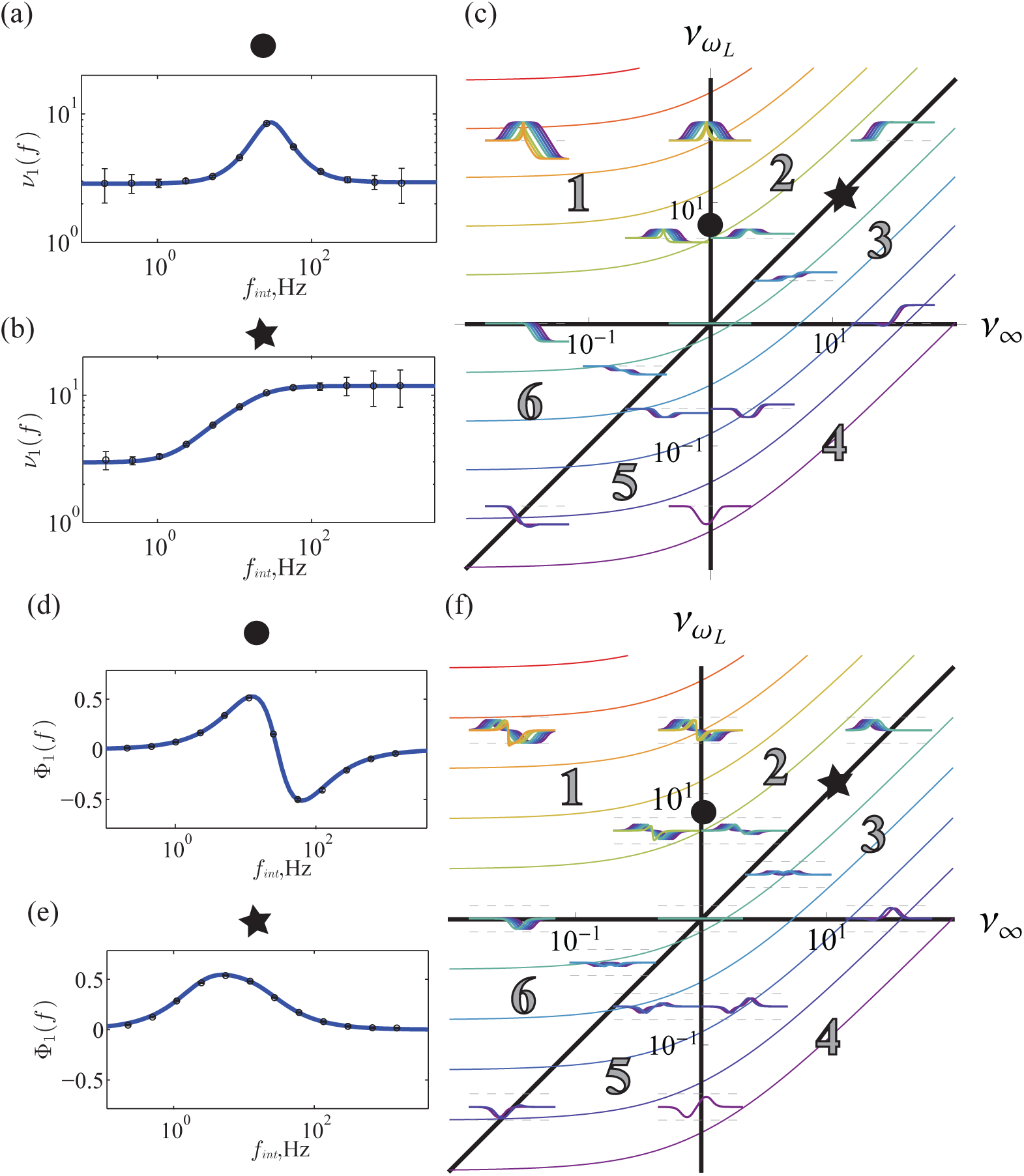
The 6 distinct filter shapes in (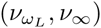)-space. (c,f) Region 1-6 denote the regions exhibiting qualitatively similar filter shapes. E.g. spiking resonance is by definition region 1 and 2. Not all of these six regions are accessible for a given *Q_L_*. Colored lines (blue to red) represent the *Q_L_*-dependent boundary below which filter shapes are forbidden because of unstable dynamics. We note that 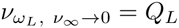. An intrinsic frequency exists in region above the *Q_L_* = 1/2 boundary. A voltage resonance exists in the region above the *Q_L_* = 1 boundary. We show the accessible subset of corresponding filter shapes at representative positions within the regions (located at 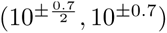 and 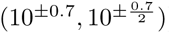) and at the border between regions (located at 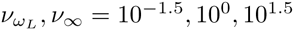). (f) Same type of plot as (c), but for the phase response. π/2 and −π/2 are shown as top and bottom bounding dashed lines for the set of phase responses at each location. The gain and phase for the position denoted by the circle are shown in (a) and (c), and for the star in (b) and (e), respectively.

In particular, depending on the region there is a peak, dip or step at *ω_L_* whose width varies with *Q_L_*. The additional high or low pass nature of the filter gives the six classes of filter shape.

While the possible shapes are simply represented in this space, the constraints are no longer represented in a plane since they depend additionally on *Q_L_*. We now dissect the effects of the stability and voltage resonance constraint on determining which filter shapes are allowed where. A main conclusion that can be drawn is that a lower bound for accessible filters is 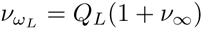 (the colored lines in Fig. 9).

With reference to Fig. 10, the stability constraint, *Q_L_* < *ω_L_τ*_w_, translates into 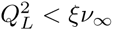 with the correct root of *ξ* given by the values of *Q_L_, v_∞_, v_low_*, and *ω_L_*.

**Figure 10.**
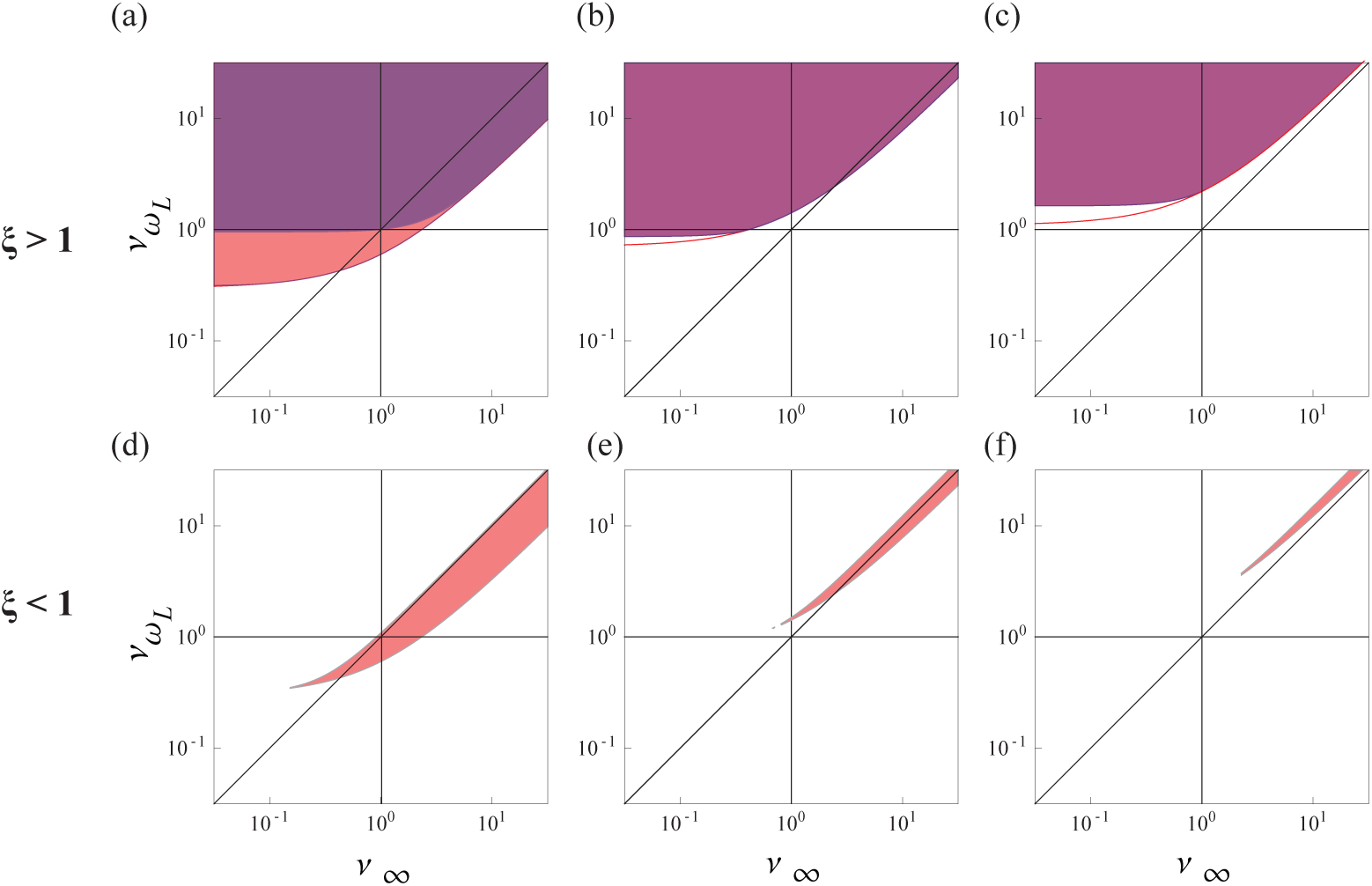
The accessible region of filter shapes depends on *Q_L_* and the relative speed of spiking to intrinsic dynamics ξ = *τ_w_*/*τ_c_*. The purple region marks the region of voltage resonant filters. This region is contained in the red region of stable filters, whose lower bound moves to larger 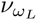 with *Q_L_*. For relatively slow intrinsic spiking (a,b,c), there are regions of non-spiking resonant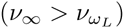, but voltage resonant filters. Filters for relatively fast intrinsic dynamics (d,e,f) only exist as high pass resonant filters for large *Q_L_*. (Left to right: *Q_L_* = 0.3, 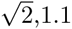. Top row: ξ = 10. Bottom row: ξ = 0.1).

Which root can also be checked by which of *τ_w_* and *τ_c_* is larger. This constraint breaks into branches when combined with the other constraints.

For ξ < 1 so that the intrinsic dynamics is faster than the spiking dynamics, the region exhibiting stable filters is constrained to a sliver, 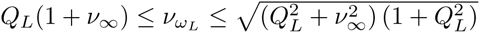, with an additional constraint on the lower bound, 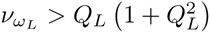 so that stable filters only exist for 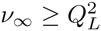 and 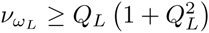 For values of *v*_∞_ and 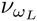 increasing from this lower bound point, the accessible region forms a band whose vertical thickness grows with *v*_∞_ and it extends out parallel with the line 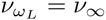 for large *v_∞_*. For increasing *Q_L_*, the accessible region shifts right and up so that the band is eventually contained in 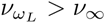 and 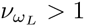 region, i.e. only high pass, resonating filters are allowed.

For ξ > 1 so that the spiking is faster than the intrinsic dynamics, the region exhibiting stable filters has no upper bound in 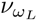. The lower bound is 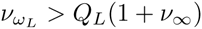 when 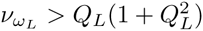 and 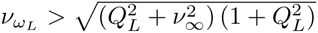 when 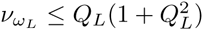 The latter bound differs significantly from *Q_L_*(1 + *v_∞_*) when *Q_L_* > 1/2.

The voltage resonance condition can also be mapped to this space by replacing *ω_L_τ_w_* by 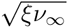 giving 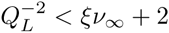. For both roots of ξ, all stable filters are voltage resonant when 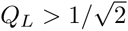

For ξ < 1, and 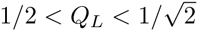 the voltage resonant filters exist at large *v_∞_* only for 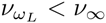, i.e. only for non-spiking resonant filters, possible because the high pass limit is brought up by the additional high pass filter above the peak of the resonance, e.g. Fig. 11. Conversely, the spiking resonant filters here lack a voltage resonance because the spiking resonance arises not from the voltage resonance but from the lower frequency amplification due to the high pass spiking filter.

**Figure 11.**
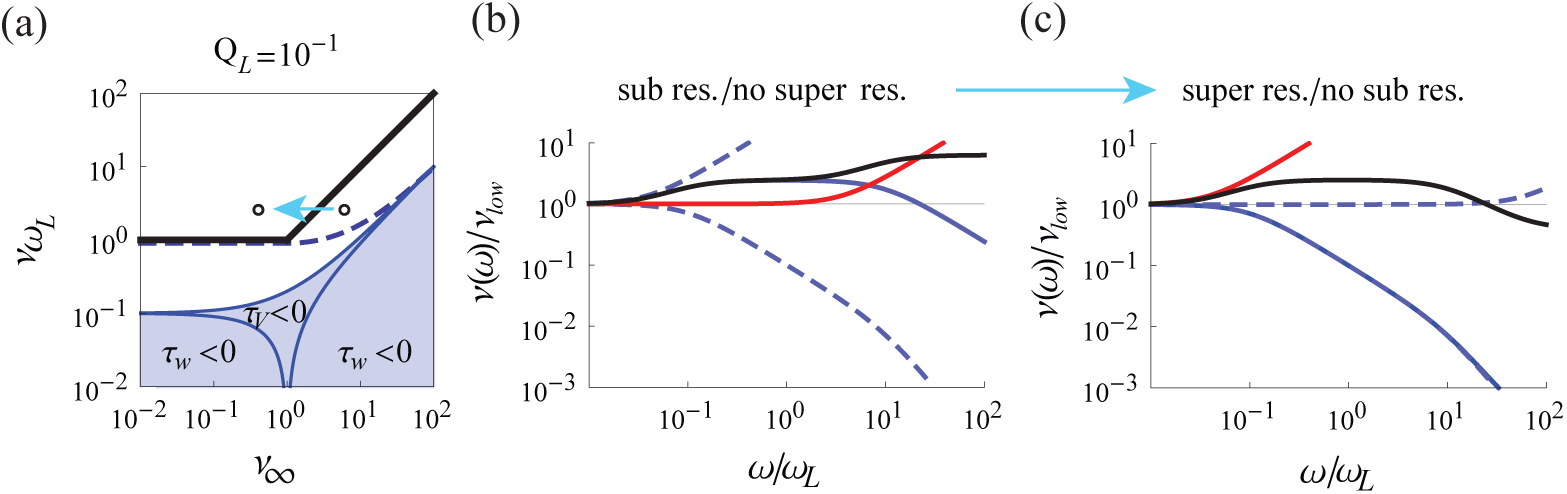
An example of filter shaping: attenuation at high frequencies uncovers an amplified band of intermediate frequencies. (a) The shape space representation showing the region of accessible filters (white) for *Q_L_* = 0.1. The blue regions exhibit unstable filters. Filters obtained from points above the thick black line are spiking resonant. Filters obtained from points above the black dashed line are voltage resonant. The arrow illustrates a path in shape space along which *v_∞_* is decreased. (b) and (c) show the beginning and end filters along the path in (a). For (b) and (c), blue dashed lines are the high and low pass components of the current-to-voltage filter, which itself is shown in solid blue. Shown in red is the voltage-to-spiking filter which combined with the current-to-voltage filter gives the full filter, shown in black.

For ξ > 1, and *Q_L_* decreasing from 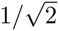, the lower bound to the voltage resonance region interpolates across *v*_∞_ from the line 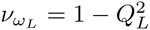, which rapidly approaches 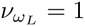 as *Q_L_* is increased, to the lower bound of the region of stable filters, 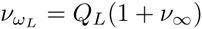 Thus, stable filters exhibit a voltage resonance when 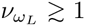, independent of *Q_L_*. The absence of a spiking resonance, 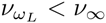, however holds over a large sub region of these stable, ξ > 1, and voltage-resonant filters, for same reason as in ξ < 1 that the high pass limit is brought up by the additional high pass filter above the peak of the resonance, thus covering it.

For *Q_L_* < 1/2, the depolarization condition, γ^−2^ < ξ*v_high_* < *γ*^2^, also excludes some regions for hyperpolarizing *w* (see Fig. 9).

The phase response across this representation is shown in Fig. 9(f). We find 0 lag when *ω* = *ω_L_* so that the input and the response are synchronous. For the spiking resonance region, we always find a delay for slower and an advance for faster input frequencies. For non-resonant cases, it is possible to observe delays or advances for both faster and slower input frequencies.

## Discussion

A neuron’s dynamic gain constrains its signal processing capabilities. Our analysis provides the first complete analysis of an expression for dynamic gain of a resonator neuron model. The level-crossing approach used here has been previously applied to 1D models to study correlation gain [23,42,44,57], dynamic response [19,37,42], and Spike-Triggered Averaged stimulus and variance [23, 42]. Consistent with conditions for the validity of the approach [19], experiments have directly demonstrated that Gauss-Rice neurons can provide a surprisingly accurate description of cortical neurons [44,58]. We find that the space of gain functions contains six types, two of which are resonant. The height of a resonant response is strictly dominated by intrinsic adaptation, while its sharpness is controlled by the strength of the subthreshold resonance. In particular, sharper peaks arise for higher intrinsic frequencies. We determined the parameter region where an intrinsic frequency exists and where subthreshold and spiking resonance are exhibited. We find that all possible combinations of the presence or absence of these three features have finite volume in parameter space. We expect profitable applications of our results to the study of the connection between intrinsic properties and population oscillations.

**Model limitations** Neuron models with hard-thresholds, such as the LIF and GIF, have been unexpectedly successful in modeling cortical neurons [58]. They are nevertheless are obtained from more complex models by a series of reductions.

In Methods, we gave a rationale for the reduction to a no-reset, hard-threshold model, where we state the additional limitations imposed by lifting the voltage reset. First, these models do not apply to mean-driven situations and so do not cover phenomena such as the masking of a subthreshold resonance by a resonance at the firing rate [29]. Second, to avoid extremely bursty spike patterns, we extend previous work [19] and argue that the correlation time of the input, *τ_I_*, and the correlation time of the voltage statistics, *τ_s_*, can not be too different. This precludes analysis involving white current noise but implies that satisfaction depends additionally on intrinsic parameters through their dependence on *τ_s_*. For example, since *τ_s_* ≤ *τ_V_*, the rough validity condition 1 ~ *τ_s_*/*τ_I_* ≲ *τ_V_*/*τ_I_* so that the timescale of the input fluctuations, *τ_I_*, should not be much slower than the membrane time constant, *τ_V_*. Third, for correspondence with threshold models the voltage relaxation time, *τ_r_*, should fall within the average inter-spike interval, *v*_0_*τ_r_* ≪ 1. Last, these models should only be considered in the irregular firing regime, *v*_0_*τ_s_* ≪ 1. We found that *τ_r_* ≤ *τ_s_* for *τ_w_* > *τ_I_*, so that this last constraint is in fact implied by the combination of the second and third.

To verify the validity of the no-reset model within the prescribed range, we made a direct, quantitative comparison to a canonical model with an active-spike generating mechanism. The dynamic gain of the two models coincides up to the high frequency limit, *flimit*, beyond which the low pass effect of the finite action potential rapidness dominates. Thus, all of the 6 distinct types of response shapes are altered by additional low pass behavior at high frequencies. For a previously used value of the rapidness, the intermediate frequency behavior is affected, while for a higher, and perhaps more accurate value it is not, and the artificially flat high frequency response is brought down by the realistic finite onset rapidness. In summary, these results show that the simplification to a no-reset, hard threshold is an adequate approximation when response features are slower than the speed of action potential onset.

A topic of related future work regards the possibility of accelerated kinetics of auxiliary currents during a spike [59]. To study such a scenario, one could numerically compute the gain for a model where the auxiliary current, *w*, undergoes a jump at spike times.

In this study of the Gauss-Rice GIF neuron and a previous on the Gauss-Rice LIF [42], exponentially-correlated Gaussian noise was used as an example of a Gaussian input statistics with non-trivial temporal correlations. These input statistics will not in general produce self-consistent firing statistics. It is therefore important to note that the approach to the linear response taken here admits arbitrary temporal correlations in the input, so long as their effect on the short-delay features of the temporal correlation of the voltage can be calculated, since that is what determines *τ_s_* and thus the effect of temporal correlations on the response properties. We also note that since the voltage correlation affects the response properties only through *τ_s_*, there is an equivalence class structure over the space of input correlation functions based on how they influence *τ_s_*.

**Relation to previous work on Type II membrane excitability** Excitable membranes are classified by the type of bifurcation that they undergo from resting to spiking, with Type I and II referring to super and sub critical Hopf bifurcation, respectively. The respective set of eigenvalues around the resting state are real and complex, with the imaginary part of the latter providing an intrinsic frequency. In this case, the voltage impulse response exhibits decaying oscillations and the voltage response function can exhibit a resonant peak near the intrinsic frequency. The mean-driven stationary spiking response rises continuously from 0 for Type I while firing in Type II neurons begins only at a finite frequency. The dynamic gain of the spiking response of Type II neurons can exhibit a superthreshold resonance arising from such subthrehsold resonance.

Frequency-sweeping ZAP input currents have revealed resonant responses from neurons in the inferior olive [60,61], thalamus [62], hippocampus [63], and cortex [64]. Consistent with the type classification, these cells often display Type II membrane excitability properties such as subthreshold oscillations with power at similar frequencies as the spiking resonance (for a review, see ref. [7]). Type II stationary spiking responses have been measured in cortical interneurons [65]. Direct measurements of the dynamic gain of resonator neurons are lacking, however. Moreover, these existing measurements used the mean input to drive the neurons to spike. Resonator response properties in the *in vivo* fluctuation-driven regime remain unmeasured.

Numerical simulations of resonator models containing the minimally required currents can nevertheless reproduce the peaked voltage and ZAP response and bimodal ISI distributions in both mean and fluctuation-driven regimes [66-68]. Inspired in part by the research presented here, Tchumatchenko and Clopath [25] used similar methods as those used here on excitatory and inhibitory GIF networks where they investigated the role of subthreshold resonance and electrical synapses on the emergence of network oscillations for a particular choice of model parameters, in which they also confirm the correspondence between the response properties with and without voltage reset. The remaining few analytical results for the stationary and linear response have so far been restricted to the long intrinsic time constant limit, *τ_w_* ≫ 1 [22,29]. In this paper, we are able to obtain exact results for the stationary and linear response for all values of *τ_w_*, something not possible in ref. [22] due to the difficulty of the analytics of the Fokker-Planck approach used there. For large *τ_w_* and the fluctuation-driven regime, our results qualitatively match their high noise results, where σ*_I_* ~ 0.1 − 1. Since Gauss-Rice models apply only to the fluctuation-driven regime, there is no meaningful mean-driven, deterministic limit attained in the limit of vanishing noise strength with which to compare to the mean-driven results of [22], such as the shift in the resonant frequency with increasing, small amounts of noise. Their phase diagram of subthreshold behavior is essentially the same as ours, up to reparametrization. We also note that the low frequency limit will differ slightly between the models due to the slightly differing slopes of their fI-curves. These small quantitative discrepancies between idealized models should not, however, be emphasized over their ability to provide a qualitative explanation of the phenomena. Finally, we note the GIF Spike-Triggered Averaged can be obtained from expression for dynamic gain. It has also been computed through other methods [69].

**Uses of the dynamical response in the theory of recurrent networks** Explicit expressions for the linear response, such as Eq. (41) obtained above, are essential ingredients for the analysis of the collective states in recurrent networks. First, they are the key quantity in the evaluation of population stability [21]. The dynamics of the population firing rate linearized around one of its fixed points is defined by the linear response function. Second, knowledge of the response function additionally reveals the correlation gain in the mapping of input current correlations to output spiking correlations. Recurrent networks exhibiting such gain will generate self-consistent patterns of inter-neuron correlations [47,49,70]. In the Gauss-Rice approach used here, the linear response providing the population stability and correlation gain is tractable for arbitrary Gaussian input current. Many networks generate such input statistics, most prominently balanced networks [71, 72]. We expect that the correlation gain and population firing rate stability of these networks can be theoretically investigated using the expressions for the linear response derived here.

One target application area is in understanding the connection between circuit oscillations and single cell excitability. Subthreshold resonance is often neglected in modeling studies of the PING and ING mechanisms for population oscillations [73]. This is despite the ample suggestive evidence of phase locking between subthreshold oscillations and gamma band population oscillations [7]. This connection has been studied in the olfactory bulb where mitral cells display a host of resonator properties such as subthreshold oscillations [5, 6], rebound spikes [74], and Type II phase resetting curves [75]. The role of this resonance in sustaining the population oscillation has not been directly assessed in detailed network models of resonating mitral cells [76], though it should play a role in either of two existing hypotheses for the origin of the oscillations [77]. Combining subthreshold and PING mechanisms has been studied in other contexts [78].

The demonstrated subthreshold resonance in inhibitory interneurons in cortex likely also contributes to the population oscillation observed there (as suggested by the numerical results of [79] and [78]) and could be investigated using the expression for dynamic gain that we provide. A first of such studies inspired by an unpublished version of the work presented here has already appeared [25], where the Gauss-Rice GIF response gain was also derived.

Finally, an *ad hoc* dynamic response filter of the same form as the one derived here [80] has been successful in modeling responses of cortical neurons (personal communication O. Shriki). The explicit dependence in our derived expression on the parameters of an underlying neuron model can be used to extend those studies, in particular, by inferring from the fitted values the properties of the intrinsic dynamics of the measured cells.

**Response properties depend on the differential correlation time** The differential correlation time, *τ_s_*, was used in a variety of ways throughout this paper.

First, it appeared in expressions for other important quantities in the theory. It appears most prominently in our expression for the fluctuation-driven voltage autocorrelation function for exponentially-correlated Gaussian input current. The result for a Type II GIF, Eq. (37), gives exponentially enveloped, oscillatory decay, with a decay constant equal to the relaxation time of the model and oscillation frequency given by the intrinsic frequency, Ω. Despite these oscillations, we find that the dynamic gain depends only on the initial falloff behavior away from 0-delay, a feature that can be shown to define, *τ_s_*. From the perspective of the response then, voltage correlation functions differ only insofar as they exhibit different *τ_s_*. The characteristic time, *τ_c_*, and thus also the attenuation of the spiking filter scales linearly with *τ_s_*, influencing the high or low pass nature of the filter accordingly.

Second, *τ_s_* appears in the validity conditions for the model. Namely, the range of valid firing rates for all Gauss-Rice neurons must lie below 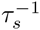.

Third, model parameters such as the intrinsic time scale, *τ_w_*, have an effect on dynamic response features, such as the high and low frequency limits, only through *τ_s_*. The analysis of their effect on *τ_s_* provides insight as to their role in sculpting the response properties. In Fig. 7(c) for example, *τ_s_* grows with *τ_w_*, and for large *τ_w_* saturates at *τ_s_*,*GIF* → *τ_s_*,*LIF* = *τ_V_*, so that *τ_s_* can only be made shorter, not longer, than the membrane time constant, *τ_V_*, by intrinsic and synaptic current parameters.

The central role of *τ_s_* could be tested by applying a variety of input correlation functions with significant differences only away from the fall-off at 0-delay so that they provide the same *τ_s_*. Our model predicts no significant change in the response properties. Such a large number of experiments could be performed by methods of high-throughput electrophysiology currently under development.

**The six filter types of the Gauss-Rice GIF** We re-expressed the response expression, Eq. (20), using the center and high frequency response relative to the low frequency response, 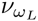 and *v_∞_* respectively. We find six qualitatively distinct filter shapes distributed around (1,1) in the 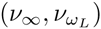 plane, with the value of *Q_L_* determining which of the six are accessible. Depending on the region there is a peak, dip or step at *ω_L_* whose width is determined by *Q_L_*. We summarize below the constraints on the accessible shapes set by *Q_L_*. For *Q_L_* < 1/2, all six filters shapes are possible for fast relative spiking (*τ_c_* < *τ_w_*). There are no high pass resonating shapes in the limit of vanishing *Q_L_* for slow relative spiking (*τ_c_* > *τ_w_*). For

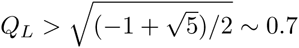 all accessible shapes have elevated response at the center frequency, 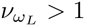. For 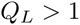, all allowed filter shapes are resonating, that is 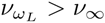. There are no low pass resonating filters for slow relative spiking and so a sharp resonance, *i.e.* a high *Q_L_*, is only possible when the overall filter is high pass.

Neither voltage nor spiking resonance strictly imply the other in this model. First, there can be voltage resonance with no spiking resonance because the spiking high pass pulls up the response in the high input frequency range above the elevated response around the intermediate-range resonant input frequency. The high frequency limitation of the approach (e.g. Fig. 12) implies that the elevated response extends up to the speed of the action potential, leaving a broad resonant band at high input frequencies. Second, there can be spiking resonance with no voltage resonance because of a low frequency attenuation by the spiking high pass filter of a low pass current-to-voltage filter.

In addition, neither voltage nor intrinsic resonance strictly imply the other. First, the existence of an intrinsic frequency does not imply voltage resonance in general because the response at *ω_L_* where Ω becomes finite is *Q_L_* = 1/2 and is thus still attenuated relative to the response at low input frequencies. This response only becomes resonant at *Q_L_* = 1. Second, there can be a voltage resonance with no intrinsic resonance for the same reason that a high pass with low characteristic frequency (this time from relatively slow intrinsic dynamics) can sculpt a peak from the low pass component of the full filter.

Finally, we found that the strength of the spiking resonance 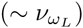 is composed of a contribution from the intrinsic timescale, *τ_w_* and from the intrinsic frequency, Ω. Nevertheless, 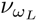 is dominated by the attenuation at low input frequencies associated with the high pass effect of large *τ_w_*, while the unique effect of Ω is to sharpen this resonance.

**The cascade representation of the dynamic response** The effect of spiking in the Gauss-Rice formulation of the response is as an explicit first-order high pass filter of the voltage dynamics (see Eq. (20)). We note that this high pass behavior associated with spiking is distinct from that discussed in the literature as arising from sodium channel inactivation [81]. This has nothing to do with the Gauss-Rice high pass arising in this paper. In this work, we always consider the threshold fixed. Closed form expressions are thus obtained for the low frequency limit and characteristic time of this filter in terms of the parameters of the model. When the characteristic frequency is high, the filter has the effect of flattening an otherwise decaying voltage response. The flattening effect is physiologically meaningful up to frequencies at which the spike-generator cut-off appears. It thus sculpts a plateau of constant response at high frequencies that can be elevated or depressed relative to the low frequency response. On the other hand, when the characteristic frequency is low, the resulting effect is a low frequency attenuation that carves out a resonant peak. The high pass characteristics are then also dependent on the intrinsic timescales.

## Methods

### Reduction from conductance-based models

Here, we detail how one arrives at a model like the one used in this paper from simplifications made to the synaptic, subthreshold, spiking, and spiking reset currents of a Hodgkin-Huxley type neuron model for the dynamics of the somatic transmembrane voltage potential, *V* (here measured in mV),

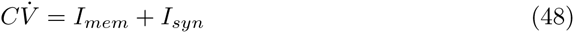

where *C* is the membrane capacitance, *I_mem_* is the sum of all membrane currents and *I_syn_* is the total synaptic current arriving at the soma. Our exposition of the reductions to synaptic and subthreshold currents is standard. To the exposition of the reductions of spiking currents we add analysis determining the high frequency limit, *f_limit_*, below which the approximation to a hard threshold is valid. To the exposition of the reduction of reset currents, we add more detailed consideration of the mechanisms through which the no reset approximation breaks down.

#### Synaptic current

*I_syn_* contributes current terms of the form *g_syn_*(*t*)(*V − E*), where *E* is the reversal potential for the synapse type and *g_syn_*(*t*) is the time-varying, synaptic input conductance for that class of synapse whose time course is determined by presynaptic activity. For a neuron embedded in a large, recurrently-connected population, this presynaptic activity arises from both the recurrent presynaptic pool of units (numbering *K ≫* 1 on average) and any external drive. In networks with sufficient dissipation, the external drive acts to maintain ongoing activity. The measured activity of networks in this regime is asynchronous and irregular and can be achieved robustly in models by an approximate 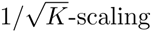 of the recurrent coupling strength, *J*. This scaling choice has the effect of balancing in the temporal average the net excitatory and inhibitory input to a cell, leaving fluctuations to drive spiking. In this *fluctuation-driven regime*, the mean-field input to a single neuron resembles a stochastic process. In the limits of (1) many, (2) weak, and (3) at most weakly correlated inputs, a diffusion approximation of *I_syn_* (*t*) can be made such that it obeys a Langevin equation [82,83]. While not yet developed for the Gauss-Rice neuron approach, analytical tools for computing the response in the case of the shot noise resulting when (1) fails are appearing [84]. Strong inputs do exist in the cortex where synaptic strengths are logarithmically distributed. Nevertheless, many strengths are weak, and we treat only (2) here. Finally, an active decorrelation in balanced networks justifies (3). Expanding *I_syn_* to leading order in the conductance fluctuations reduces the input to additive noise yielding the Gaussian approximation to the voltage dynamics, also known as the effective time constant approximation [84,85]. The quality of this approximation depends on the relative difference between the reversal potential and the voltage. Somas receive input from two broad classes of synapse: excitatory ones for which the difference is large, and inhibitory ones for which the difference is smaller so that they are less well-approximated. The two types can also differ in their kinetics. While both are generally low pass, their characteristic times can be different. Their combination can thus have qualitative effects on the response [19]. We retain only a single synapse type so as to concentrate on the shaping of the filter properties by the intrinsic currents of the neuron model.

In this approximation to additive Gaussian noise, the time-dependent ensemble from which the input signal, *I_syn_*(*t*), is sampled is completely described by a *variance channel* carrying the dynamics of the fluctuations of the network activity, and a *mean channel* carrying the dynamics of the mean network activity. More complicated compound input processes described by higher order statistics offer more channels but they are negated by the diffusion approximation to a Gaussian process. The variance channel determines the fluctuations of *I_syn_* (t) on which rides a DC component described by the mean channel. We can thus write

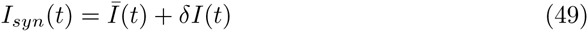

where the zero-mean Gaussian process *δI*/(*t*) is characterized by the variance, 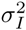, and correlation time, τ*_I_*, of the fluctuations, both of which can in general vary in time, and *Ī*(*t*) is the time-dependent population mean. The population mean of a quantity, *x*, will be denoted by a bar so that 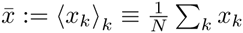, where *k* indexes the neuron. For stationary input, the time average of 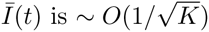 due to the balance. In this paper, we consider deterministic changes in the mean channel, *Ī*(*t*), produced for example by a global and time-dependent external drive. We compute the resulting frequency and phase response, and leave the analysis of the variance channel to a forthcoming work. For much of the paper, we will also remove explicit dependence of the model’s behavior on the input by setting σ*_I_* for a desired output firing rate and measuring time relative to *τ_I_*.

#### Subthreshold current

In the most simple case (no longer exactly the Hodgkin-Huxley formalism), each somatic current, *I_mem_,_i_,* contributes additively to *I_mem_* with a term of the form

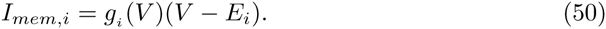

where *g_i_*(*V*) is a voltage-dependent conductance, whose effect on the voltage dynamics depends on the driving force, *V* − *E_i_*, the difference of the voltage and the reversal potential, *E_i_*. *g_i_* obeys kinetic equations based on channel activation whose specification is often made *ad hoc* to fit the data since the details of the conformational states and transitions of a neuron’s ion channels is often unknown or at least not yet well understood. Nevertheless, for voltages below the threshold for action potential initiation the voltage dynamics can be well-approximated by neglecting spike-generating currents and linearizing the dynamics of the subthreshold gating variables around the resting potential, *V\**. Following ref. [29], the resulting subthreshold dynamics is then given by

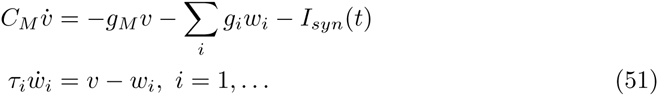

where *v* = *V − V*\* and 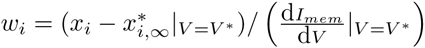 are the linearized variables for the voltage and subthreshold gating variable, *x_i_*, respectively; 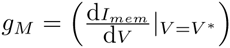 and 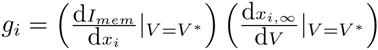 are the effective membrane conductances for the leak and for *x_i_*, respectively; and *τ_i_* = *τ_i_*(*V*\*) is the time constant of the dynamics of *w_i_*. *C_M_* is the capacitance of the membrane. The *w* variables have dimensions of voltage. Activation and inactivation gating variables have *g_i_* > 0 and *g_i_* < 0, respectively. We denote the linearized voltage by *V* instead of *v* throughout the paper to better distinguish it from the firing rate, *v*.

With the addition of a hard (i.e. sharp and fixed) voltage threshold and a reset rule to define the spiking dynamics, this defines the GIF class of models [29]. Among the models considered in ref. [29], the simplest has only one additional degree of freedom,

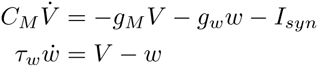

with spikes occurring at upward crossings of the threshold, *θ*. With time in units of *τ_w_* the authors multiply the voltage equation by *τ_w_*/*C_M_* and analyze the behavior as a function of two dimensionless model parameters, *α* = *g_M_τ_w_*/*C_M_* and *β* = *g_w_τ_w_*/*C_M_*, upon which the qualitative shape of the current-to-voltage filter for white noise input depends.

We consider correlated noise input that introduces an additional time scale which serves as a more natural time unit. We are also interested in the explicit dependence on *τ_w_*. Thus, we retain both of the timescales of the neuron model, *τ_V_* and *τ_w_*. We then parametrize our model using the relative conductance *g* = *β/α* = *g_w_/g_M_*, the relative membrane time constant *τ_V_*/*τ_I_* = α^−1^ = *C_M_*/*τ_I_g_M_*, and the relative *w* timescale, *τ_w_*/*τ_I_* = *β*/*τ_Ig_*. Input variance is independently fixed in order to achieve a desired firing rate. We thus make a slight alteration to the model in ref. [29],

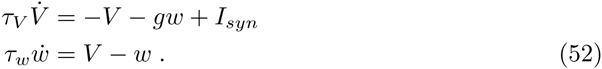

We have absorbed the 1/*g_M_* factor into the units of *I_syn_* so that all dynamic quantities are in dimensions of voltage. We keep *τ_V_* > 0 by setting *g_M_* > 0, that together with *g* > −1, this gives stable voltage dynamics. This model is the same as the one stated at the beginning of the Results section, Eq. (1).

The approximation to a hard threshold from a set of spike-generating currents that are in principle contained in *I_mem_* but are not considered explicitly in [29] involves some assumptions and approximations that have since been nicely formalized in [24] and so we include them in the following section.

#### Spike-activating current

The formulation of spike-activating currents can be simplified using the fact that all the information that the neuron provides to downstream neurons is contained in the times of its action potentials and not their shape. Only the voltage dynamics contributing to this time is retained in the model; namely, the summed rise of voltage-gated activation of the spike-generating *x_i_*, summed into a single function, *ψ*(*V*), dependent only on the voltage when its dynamics is relatively fast [24]. *ψ*(*V*) then appears as a term in the voltage dynamics and, when supralinear in *V*, acts as the spike-generating instability that, in the absence of superthreshold, hyperpolarizing currents, causes the voltage to diverge in finite time. These latter currents are simply omitted and the time at which the voltage has diverged is used in these models as the spike time. The socalled spike slope factor [24], Δ*_T_*, is the inverse curvature of the I-V curve near threshold and sets the slope of the rise of the action potential, with smaller values giving steeper rise. Its value should be measured at the site of action potential initiation, the precise location of which is not yet known in general. An upper bound on the realistic range of Δ*_T_* is, however, likely smaller than that achievable by conventional Hodgkin-Huxley-like models, even with multiple compartments [37,39], and this speed has motivated neuron models with fast action potential onset rapidness [86].

The time between the crossing of a fixed threshold voltage, *V_T_*, defined implicitly by 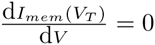, and the spike time vanishes quickly with 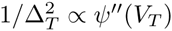, so that the further approximation to a hard threshold, i.e. for omitting *ψ*(*V*) altogether by setting the spike time at *V_T_*, becomes good for Δ*_T_* → 0. However, the instantaneous rise in voltage in this limiting approximation introduces artefactually fast population responses at high input frequencies, denoted by *f*, raising the scaling behavior to 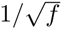 and constant for white and colored noise, respectively [20]. Nevertheless, since the discrepancy begins above some *f_limit_* depending on Δ*_T_*, the artefact can be safely ignored by considering the shape of the response only for *f* < *f_limit_*. Conveniently, an upper bound on realistic values of Δ*_T_* given by the surprisingly quick rise of real action potentials leads to a value of *f_limit_* well beyond the range of input frequencies over which realistic filtering timescales act. As a result, the approximation to a hard threshold does not alter the sub-spiking timescale response properties of the full model.

For concreteness, a popular choice for *ψ*(*V*) is *ψ*(*V*) = exp [(*V − V_T_*) /Δ*_T_*], the family of so-called exponential integrate-and-fire (EIF) models [87] for which the difference between the threshold crossing and the spike time vanishes very fast as 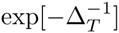. Its high frequency response falls off as 1/*f*, with a high frequency cut-off 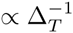. We consider an EIF version of our model defined having an additional, superlinear term in the 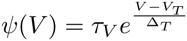. We note that a similar comparison is made in [25]. The approximate upper limit of input frequencies, *f_limit_*, below which the no-reset approximation is valid is given implicitly by the intersection of the response of the simplified model computed in this paper and the analytical high frequency response of the EIF version of the full model, computed from an expansion of the corresponding Fokker-Planck equation in *ω*^−1^ = 1/(2π*f*). We choose examples where the intrinsic dynamics are slow relative to the cut-off so we use the high frequency limit result of the EIF with no additional degree of freedom calculated in [24],

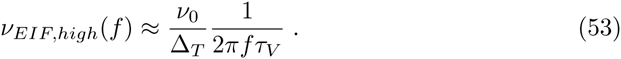

The high frequency limit of the Gauss-Rice GIF is Eq. (43). Equating these two expressions, we obtain

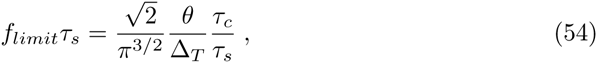

where *g*, *τ_c_*, and *τ_s_* are parameters defined later. We check this condition through numerical simulations of the EIF-version of the model. Instead of the heuristic constraints for choosing the integration time step d*t* as specified in [24], we more simply obtain the *f^−^*^1^ fall-off by raising the numerical voltage threshold for spiking, allowing the speed of the action potential to play a role at higher frequencies. While this gives an artifact in the phase response (not shown), the high frequency limit of the gain is correct. Two example gain functions are shown in Fig. 12 for a widely used value of Δ*_T_* = 3.5mV(0.35 in our units), and a value an order of magnitude smaller, Δ*_T_* = 0.35mV(0.035 in our units). The former value gives a cut-off slow enough that it affects the resonant feature, while the latter value gives a cut-off high enough that it does not. The features of the filter in this case are thus well below *f_limit_*.

**Figure 12.**
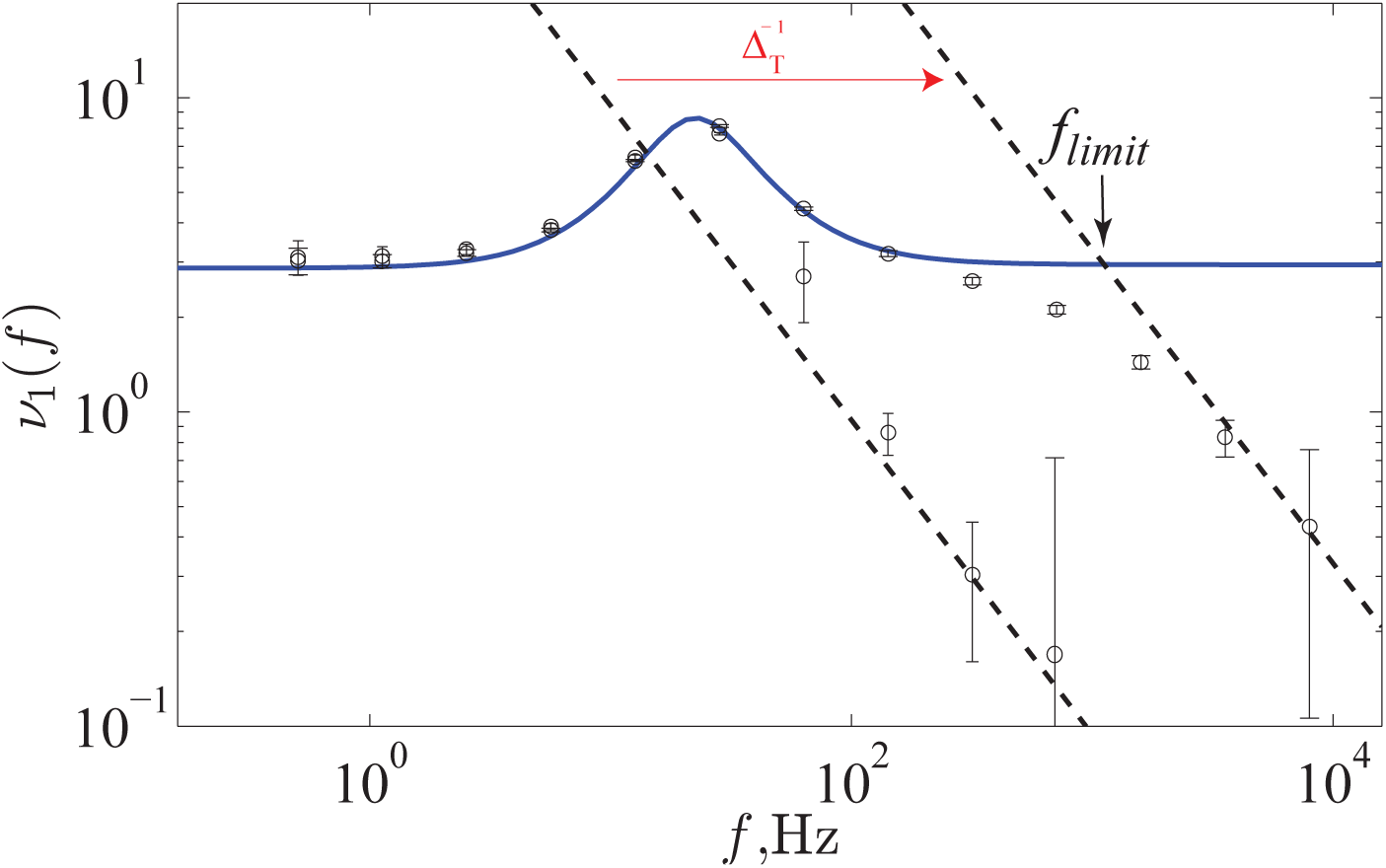
Correspondence of response between analytical result of no-reset model (blue line) and the numerical result of its EIF version (black circles). The correspondence holds up to a high frequency cut-off, *f_limit_* (Eq. (54)), due to finite rise time of action potential controlled by Δ*_T_* = 0.35,0.035. The EIF-version was simulated with *V_thr_* = 1.15, 3, and *V_T_* = 0.8, −1 (the latter was adjusted to keep *v*_0_ = 2Hz fixed). The black dashed lines correspond to the high frequency limit of the response of the EIF-type model (Eq. (53)). The no reset model had the default parameters.

Notably, the LIF FP methods have been used to obtain the linear response to a piecewise linear models [33, 34]. In these works, the high frequency artifacts induced by the hard threshold are treated explicitly and removed.

#### Resetting current

Models that neglect the downward part of the action potential require the addition of, or have already built-in a reset voltage to which the voltage is reset after a spike. The reset makes the dynamics discontinuous and a closed form expression for the frequency response for more-than-1D models appear intractable. We forgo this reset rule in order to open up the problem for deeper analysis. With this simplification, however, come three issues that we avoid by narrowing the scope of the analysis.

First, without the reset and for the case of mean-driven activity, the mean voltage is taken into an unrealistic, super-threshold range. Thus, only fluctuation-driven activity with low, subthreshold mean input is covered by this approximation, leaving out mean-driven phenomena such as the masking of a subthreshold resonance by a resonance at the firing rate as shown, e.g. in ref. [29]. This is nevertheless the operating regime of cortical networks that we wish to study. We thus set the mean input to 0.

Second, the lack of reset produces periods of artefactually high and low firing rates for respectively small and large values of the input correlation time, *τ_I_*, relative to the voltage correlation time defined here as the *differential correlation time*, 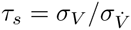. *τ_s_* is the quadratic approximation to the voltage correlation function around 0-delay (discussed in detail in the main text). This definition precludes the use of white noise input whose correlation function is non-differentiable around 0-delay. Indeed, the fractal nature of the voltage traces when the no-reset model is driven by white noise endows the model with the problematic feature that every threshold crossing has in its neighborhood infinitely many such crossings [57]. A version of this effect explains the discrepancy between reset and non-reset dynamics even in the finite realm where *τ_I_*/*τ_s_* ≪ 1. In the other limit, *τ_I_*/*τ_s_* ≫ 1 means that the voltage stays super threshold for long spans of time and so must also be excluded. Badel compares the stationary response of the LIF with and without reset across *τ_I_*, finding correspondence only in a fairly tight band around the membrane time constant, *τ_V_*, from *τ_I_* = 0.5*τ_V_* to *τ_I_* = 2*τ_V_* [19]. Given that the stationary response of the LIF also deviates from more realistic models, in this paper we do not aim for exact correspondence with the LIF but rather analyze the more general and less strong condition, *τ_I_*/*τ_s_* ~ 1, which reduces to a less strong version of the one Badel used for the LIF where 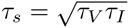. From the derivation of *τ_s_* for the Gauss-Rice GIF exposed in the main text, the condition *τ_I_*/*τ_s_* ~ 1 implies that the membrane time constant is no longer required to lie within an order of magnitude of *τ_I_* but that the validity now holds around a manifold in the space of intrinsic parameters of the model.

Third, for those neurons that do exhibit reset-like dynamics, this approach can nevertheless provide a good approximation so long as the model dynamics allow for the sample paths of the voltage trajectory after a spike with and without reset to converge onto one another before the next spike occurs. The formal condition for this is *v*_0_*τ_r_* ≪ 1, where *v*_0_ is the firing rate and *τ_r_* is the relaxation time of the deterministic dynamics of the voltage, i.e. the negative of the largest real part of the eigenvalues of the solution to the linearized voltage dynamics. For the case of 2D linear dynamics considered in this paper, with differential matrix operator *B*, 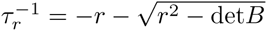 when r^2^ > det *B* and 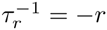 where *r* is the real part of the complex eigenvalue (see next paragraph for details). For relatively fast intrinsic kinetics, this constraint limits the range of parameters and output firing rates over which the no-reset model approximates reset dynamics to within some tolerance. However, we will show that, for relatively slow intrinsic kinetics, the condition *τ_r_* ≲ *τ_s_* holds up to a saturation level, and this together with *v*_0_*τ_s_* ≪ 1 (a condition that all healthy Gauss-Rice neurons must satisfy) guarantees the near equivalence of reset and no-reset dynamics, independent of the other parameters. In other words, the approximation is valid in this regime if the relaxation time falls within a correlated window of voltage trajectory as this is a lower bound to the time between spikes. Indeed, for any temporally correlated dynamics, it always takes some time for the state to move some fixed amount. In this context, that effect induces an relative refractory period in reset dynamics as the state must move from reset to threshold again in order to spike. It is not absolute because this time depends on the firing rate. The same type of refractoriness emerges in non-reset dynamics as the voltage must fall back below threshold in order to pass it from below again.

### Parametrization of the model

When the eigenvalues of the solution to the voltage dynamics are complex, we can re-express the denominator of Eq. (25) using the intrinsic frequency, i.e. the imaginary part of the eigenvalues of the voltage solution. We first obtain the eigenvalues. For the linear matrix evolution operator

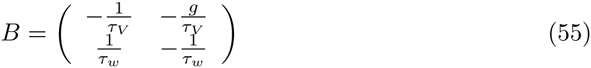

the eigenvalues are obtained via the identity

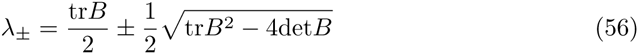

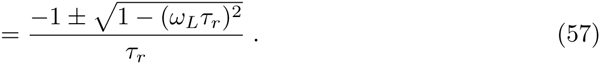

where 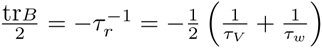 as the negative reciprocal of the harmonic mean of the two time constants, *τ_r_*, and 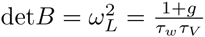 where *ω_L_* is the center frequency of the voltage filter. When *ω_L_τ_r_* < 1, the magnitude 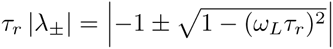.

When *ω_L_τ_r_* > 1, the eigenvalues are complex with 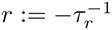 as the real part. We define the imaginary part that plays the role of the intrinsic frequency, Ω > 0, via λ_±_ = *r* ± *iΩ*, so 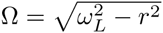 and now the magnitude is 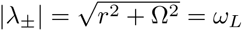. We can substitute the expression for *ω_L_*, obtaining the relation between *g* and Ω, Eq. (3),

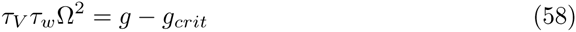

where 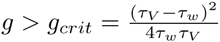 is the condition for complex eigenvalues (see Fig. 1).

### Obtaining the response function directly from spike times

Here we rederive the linear relationship between the vector strength and the linear response. *v*_1_(*ω*) from Eq. (9) can be expressed using the complex response vector,

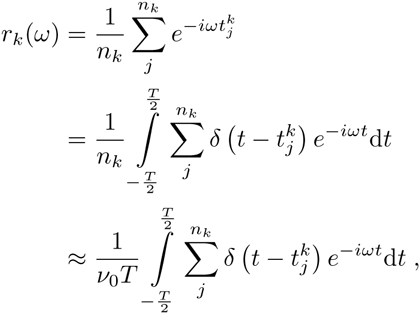

where in the last step we use *n_k_ ≈ v_0_T*, good when *T* is made much larger than 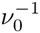. Taking the ensemble average,

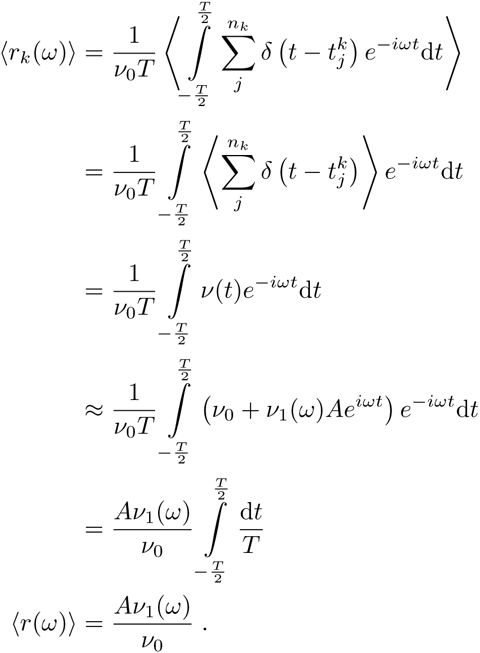

Using the decomposition of the response into its gain and phase, *v*_1_(*ω*) = |*v*_1_(*ω*)| *e*^iΦ(^*^ω^*^)^, the dynamic gain is thus obtained from the norm of the ensemble-averaged response vector, called the vector strength,

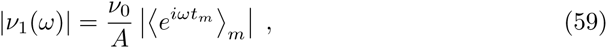

where here we have simplified the notation by having *m* run over all the spikes from the full ensemble. We computed this expression using the spike times obtained directly from numerical simulations of the stochastic dynamics generated by the neuron model. We use the result to confirm the validity of the analytical gain function derived below, whose utility goes far beyond the numerical result because it provides the explicit dependence on the model parameters.

### Computing the input variance for given firing rate

Rearranging the expression for *v*_0_ and then substituting in the *σ_I_*-dependent expression for the voltage fluctuations, *σ*, we have

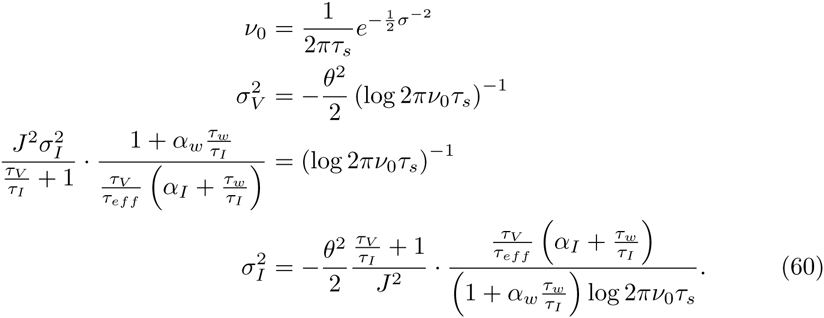

When we study the model’s behavior we will use this relation to set the input variance for a chosen output firing rate so that the dimensions of the parameter space to be explored are the four time scales in the problem, (*τ_V_*, *τ_eff_*, *τ_w_*, 1/*v*_0_) and when Ω exists, (*τ_V_*, 1/Ω, *τ_w_*, 1/*v*_0_).

### Step response

The firing rate response derived in this paper allows us to compute the response to any weak signal and we demonstrate that in this section where we derive the response to step-like input. The time-domain version of linear frequency response, *v*_1_(*t*), is the impulse response function, which when convolved with any input times series gives the corresponding response time series,

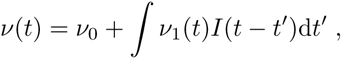

where 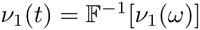 has units of [*Time*]^−2^ [*Current*]^−1^. If there is an accessible frequency representation of the input, the interaction can be made in the frequency domain and then the result transformed back to the time domain,

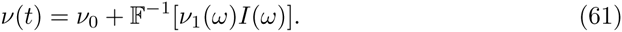

We used this definition to study the response to step-like input, *I*(*t*) = *A*Θ(*t*), with step height, *A*, and with frequency domain expression for the Heaviside theta function,

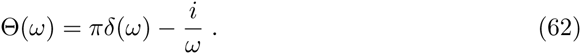

Applying the inverse Fourier transform to the product of this with the linear frequency response gives the expression for the response. The relative response is then,

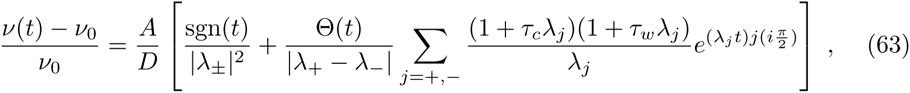

with 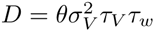. We can express Eq. (63) in terms of *r* and Ω,

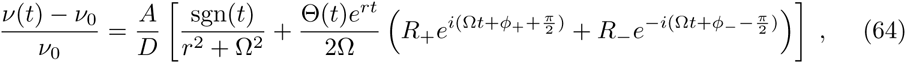

where 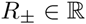 and 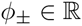 depend on the parameters. Taking the limit *t* → 0^+^, the relative instantaneous jump height is 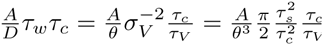, consistent with the notion that higher characteristic cutoff frequencies, *i.e.* 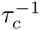, imply stronger instantaneous transmission. The exponent of the subsequent decay is 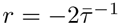, providing an envelope that funnels into the relative asymptotic response, 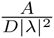, attained in the limit t → ∞. Since the oscillation amplitude scales as 1/Ω while the asymptotic response scales with 1/Ω^2^, there will be a tapering envelope for Ω > 1. Within this envelope the response oscillates at the intrinsic frequency and with a phase that is explicitly dependent on the neuron parameters, as well as implicitly though *τ_c_*. This function was used to calculate the step response shown in Fig. 2.

## Acknowledgments

MPT would like to acknowledge discussions with Tatjana Tchumatchenko at the outset of this research. FW and MPT would like to acknowledge interactions with Mike Gutnick, Andreas Neef, Elinor Lazarov, Nicolas Brunel, Srdjan Ostojic, and Ricardo Merino. We acknowledge the financial support by the Bundesministerium fur Bildung und Forschung (http://www.bmbf.de, 01GQ1005B, 01GQ07113), German-Israeli Foundation (http://www.gif.org.il, I-906-17.1/2006), Volkswagenstiftung Stiftung (http://www.volkswagenstiftung.de, ZN2632), Deutsche Forschungsgemeinschaft (http://www.dfg.de/index.jsp, SFB889), and the Max Planck Society (www.mpg.de). The funders had no role in study design, data collection and analysis, decision to publish, or preparation of the manuscript.

## References

1 Hodgkin AL. The local electric changes associated with repetitive action in a non-medullated axon. Journal of Physiology. 1948;107(2):165–181.

2 Rinzel J, Ermentrout GB. Analysis of neural excitability and oscillations. In: Koch C, Segev I, editors. Methods in Neuronal Modeling; 1998.

3 Shadlen MN, Newsome WT. The variable discharge of cortical neurons: implications for connectivity, computation, and information coding. Journal of Neuroscience. 1998 May;18(10):3870–3896. Available from: http://www.ncbi.nlm.nih.gov/pubmed/9570816.

4 Balu R, Larimer P, Strowbridge BW. Phasic stimuli evoke precisely timed spikes in intermittently discharging mitral cells. Journal of Neurophysiology. 2004 Aug;92(2):743–53. Available from: http://www.ncbi.nlm.nih.gov/pubmed/15277594.

5 Desmaisons D, Vincent JD, Lledo PM. Control of action potential timing by intrinsic subthreshold oscillations in olfactory bulb output neurons. Journal of Neuroscience. 1999 Dec;19(24):10727–37. Available from: http://www.ncbi.nlm.nih.gov/pubmed/10594056.

6 Lepousez G, Lledo PM. Odor Discrimination Requires Proper Olfactory Fast Oscillations in Awake Mice. Neuron. 2013;80(4):1010–1024. Available from: http://dx.doi.org/10.1016/j.neuron.2013.07.025.

7 Hutcheon B, Yarom Y. Resonance, oscillation and the intrinsic frequency preferences of neurons. Trends in Neurosciences. 2000;23(5):216–222.

8 Knight BW. Dynamics of encoding in a population of neurons. The Journal of General Physiology. 1972;59:734–766.

9 Carandini M, Mechler F. Spike train encoding by regular-spiking cells of the visual cortex. Journal of Neurophysiology. 1996;76(5):3425. Available from: http://jn.physiology.org/content/jn/76/5Z3425.full.pdf.

10 Higgs MH, Spain WJ. Conditional bursting enhances resonant firing in neocortical layer 2-3 pyramidal neurons. Journal of Neuroscience. 2009 Feb;29(5):1285–99. Available from: http://www.ncbi.nlm.nih.gov/pubmed/19193876.

11 Köndgen H, Geisler C, Fusi S, Wang XJ, Lüscher HR, Giugliano M. The dynamical response properties of neocortical neurons to temporally modulated noisy inputs in vitro. Cerebral Cortex. 2008 Sep;18(9):2086–97. Available from: http://www.pubmedcentral.nih.gov/articlerender.fcgi?artid=3140196&tool=pmcentrez&rendertype=abstract.

12 Boucsein C, Tetzlaff T, Meier R, Aertsen A, Naundorf B. Dynamical response properties of neocortical neuron ensembles: multiplicative versus additive noise. Journal of Neuroscience. 2009 Jan;29(4):1006–10. Available from: http://www.ncbi.nlm.nih.gov/pubmed/19176809.

13 Silberberg G, Bethge M, Markram H, Pawelzik K, Tsodyks M. Dynamics of population rate codes in ensembles of neocortical neurons. Journal of Neurophysiology. 2004 Feb;91(2):704–9. Available from: http://www.ncbi.nlm.nih.gov/pubmed/14762148.

14 Pozzorini C, Naud R, Mensi S, Gerstner W. Temporal whitening by power-law adaptation in neocortical neurons. Nature Neuroscience. 2013;16(7):942–8. Available from: http://www.ncbi.nlm.nih.gov/pubmed/23749146.

15 Lundstrom BN, Higgs MH, Spain WJ, Fairhall AL. Fractional differentiation by neocortical pyramidal neurons. Nature Neuroscience. 2008;11(11):1335–1342.

16 Testa-Silva G, Verhoog MB, Linaro D, de Kock CPJ, Baayen JC, Meredith RM, et al. High Bandwidth Synaptic Communication and Frequency Tracking in Human Neocortex. PLoS Biology. 2014;12(11):e1002007. Available from: http://dx.plos.org/10.1371/journal.pbio.1002007.

17 Volgushev M, Ilin V, Stevenson IH. Identifying and Tracking Simulated Synaptic Inputs from Neuronal Firing: Insights from In Vitro Experiments. PLOS Computational Biology. 2015;11(3):e1004167. Available from: http://dx.plos.org/10.1371/journal.pcbi.1004167.

18 Ostojic S, Szapiro G, Schwartz E, Barbour B, Brunel N, Hakim V. Neuronal Morphology Generates High-Frequency Firing Resonance. Journal of Neuroscience. 2015;35(18):7056–7068. Available from: http://www.jneurosci.org/cgi/doi/10.1523/JNEUR0SCI.3924-14.2015.

19 Badel L. Firing statistics and correlations in spiking neurons: A level-crossing approach. Physical Review E. 2011 Oct;84(4):041919. Available from: http://link.aps.org/doi/10.1103/PhysRevE.84.041919.

20 Brunel N, Chance F, Fourcaud N, Abbott L. Effects of Synaptic Noise and Filtering on the Frequency Response of Spiking Neurons. Physical Review Letters. 2001 Mar;86(10):2186–2189. Available from: http://link.aps.org/doi/10.1103/PhysRevLett.86.2186.

21 Brunel N, Hakim V. Fast global oscillations in networks of integrate-and-fire neurons with low firing rates. Neural Computation. 1999 Oct;11(7):1621–71. Available from: http://www.ncbi.nlm.nih.gov/pubmed/10490941.

22 Brunel N, Hakim V, Richardson M. Firing-rate resonance in a generalized integrate-and-fire neuron with subthreshold resonance. Physical Review E. 2003 May;67(5). Available from: http://link.aps.org/doi/10.1103/PhysRevE.67.051916.

23 Burak Y, Lewallen S, Sompolinsky H. Stimulus-dependent correlations in threshold-crossing spiking neurons. Neural Computation. 2009;2308:2269–2308. Available from: http://www.mitpressjournals.org/doi/abs/10.1162/neco.2009.07-08-830.

24 Fourcaud-Trocmé N, Hansel D, Van Vreeswijk C, Brunel N. How spike generation mechanisms determine the neuronal response to fluctuating inputs. Journal of Neuroscience. 2003 Dec;23(37):11628–40. Available from: http://www.ncbi.nlm.nih.gov/pubmed/14684865.

25 Tchumatchenko T, Clopath C. Oscillations emerging from noise-driven steady state in networks with electrical synapses and subthreshold resonance. Nature Communications. 2014;5:1–9. Available from: http://dx.doi.org/10.1038/ncomms6512.

26 Geisler C, Brunel N, Wang XJ. Contributions of intrinsic membrane dynamics to fast network oscillations with irregular neuronal discharges. Journal of Neurophysiology. 2005 Dec;94(6):4344–61. Available from: http://www.ncbi.nlm.nih.gov/pubmed/16093332.

27 Middleton JW, Chacron M, Lindner B, Longtin A. Firing statistics of a neuron model driven by long-range correlated noise. Physical Review E. 2003;68(February):1–8.

28 Ostojic S, Brunel N, Hakim V. How connectivity, background activity, and synaptic properties shape the cross-correlation between spike trains. Journal of Neuroscience. 2009;29(33):10234–53. Available from: http://www.ncbi.nlm.nih.gov/pubmed/19692598.

29 Richardson MJE, Brunel N, Hakim V. From subthreshold to firing-rate resonance. Journal of Neurophysiology. 2003 May;89(5):2538–54. Available from: http://www.ncbi.nlm.nih.gov/pubmed/12611957.

30 Lindner B, Schimansky-Geier L. Transmission of noise coded versus additive signals through a neuronal ensemble. Physical Review Letters. 2001;86(14):2934–2937.

31 Bernardi D, Lindner B. A frequency-resolved mutual information rate and its application to neural systems. Journal of Neurophysiology. 2015;113(5):1342–1357. Available from: http://jn.physiology.org/lookup/doi/10.1152/jn.00354.2014.

32 Eyal G, Mansvelder HD, de Kock CPJ, Segev I. Dendrites impact the encoding capabilities of the axon. Journal of Neuroscience. 2014;34(24):8063–71. Available from: http://www.ncbi.nlm.nih.gov/pubmed/24920612.

33 Wei W, Wolf F. Spike Onset Dynamics and Response Speed in Neuronal Populations. Physical Review Letters. 2011 Feb;106(8):1–4. Available from: http://link.aps.org/doi/10.1103/PhysRevLett.106.088102.

34 Wei W, Wolf F, Wang Xj. Impact of membrane bistability on dynamical response of neuronal populations;.

35 Naundorf B, Geisel T, Wolf F. Dynamical response properties of a canonical model for type-I membranes. Neurocomputing. 2005 Jun;65–66:421–428. Available from: http://linkinghub.elsevier.com/retrieve/pii/S0925231204004254.

36 Naundorf B, Geisel T, Wolf F. Action potential onset dynamics and the response speed of neuronal populations. Journal of Computational Neuroscience. 2005;18(3):297–309.

37 Naundorf B, Wolf F, Volgushev M. Unique features of action potential initiation in cortical neurons. Nature. 2006 Apr;440(7087):1060–1063. Available from: http://www.ncbi.nlm.nih.gov/pubmed/16625198.

38 Gerstner W. Population dynamics of spiking neurons: fast transients, asynchronous states, and locking. Neural Computation. 2000;12(1):43–89.

39 Ilin V, Malyshev A, Wolf F, Volgushev M. Fast computations in cortical ensembles require rapid initiation of action potentials. Journal of Neuroscience. 2013 Feb;33(6):2281–92. Available from: http://www.ncbi.nlm.nih.gov/pubmed/23392659.

40 Huang M, Volgushev M, Wolf F. A small fraction of strongly cooperative sodium channels boosts neuronal encoding of high frequencies. PLoS ONE. 2012;7(5).

41 de Solages C, Szapiro G, Brunel N, Hakim V, Isope P, Buisseret P, et al. High-Frequency Organization and Synchrony of Activity in the Purkinje Cell Layer of the Cerebellum. Neuron. 2008;58(5):775–788.

42 Tchumatchenko T, Wolf F. Representation of Dynamical Stimuli in Populations of Threshold Neurons. PLoS Computational Biology. 2011 Oct;7(10):e1002239. Available from: http://dx.plos.org/10.1371/journal.pcbi.1002239.

43 Tchumatchenko T, Malyshev A, Wolf F, Volgushev M. Ultrafast Population Encoding by Cortical Neurons. Journal of Neuroscience. 2011 Aug;31(34):12171–12179. Available from: http://www.jneurosci.org/cgi/doi/10.1523/JNEUR0SCI.2182-11.2011.

44 Tchumatchenko T, Malyshev A, Geisel T, Volgushev M, Wolf F. Correlations and Synchrony in Threshold Neuron Models. Physical Review Letters. 2010 Feb;104(5):5–8. Available from: http://link.aps.org/doi/10.1103/PhysRevLett.104.058102.

45 Shea-Brown E, Josić K, de la Rocha J, Doiron B. Correlation and Synchrony Transfer in Integrate-and-Fire Neurons: Basic Properties and Consequences for Coding. Physical Review Letters. 2008 Mar;100(10):1–4. Available from: http://link.aps.org/doi/10.1103/PhysRevLett.100.108102.

46 de la Rocha J, Doiron B, Shea-Brown E, Josić K, Reyes A. Correlation between neural spike trains increases with firing rate. Nature. 2007 Aug;448(7155):802–6. Available from: http://www.ncbi.nlm.nih.gov/pubmed/17700699.

47 Hu Y, Trousdale J. Motif statistics and spike correlations in neuronal networks. Journal of Statistical Mechaniics. 2013;03012. Available from: http://iopscience.iop.org/1742-5468/2013/03/P03012.

48 Vilela RD, Lindner B. Comparative study of different integrate-and-fire neurons: Spontaneous activity, dynamical response, and stimulus-induced correlation. Physical Review E. 2009 Sep;80(3):031909. Available from: http://link.aps.org/doi/10.1103/PhysRevE.80.031909.

49 Trousdale J, Hu Y, Shea-Brown E, Josić K. Impact of network structure and cellular response on spike time correlations. PLoS Computational Biology. 2012 Jan;8(3):e1002408. Available from: http://www.pubmedcentral.nih.gov/articlerender.fcgi?artid=3310711&tool=pmcentrez&rendertype=abstract.

50 Rosenbaum R, Marpeau F, Ma J, Barua A, Josić K. Finite volume and asymptotic methods for stochastic neuron models with correlated inputs. Journal of Mathematical Biology. 2012;65(1):1–34.

51 Young G. Note on excitation theories. Psychometrika. 1937 Jun;2(2):103–106. Available from: http://www.springerlink.com/index/10.1007/BF02288064.

52 Izhikevich EM. Resonate-and-fire neurons. Neural Networks. 2001;14(6–7):883–894. Available from: http://www.ncbi.nlm.nih.gov/pubmed/18244602.

53 Schleimer JH, Stemmler M. Coding of Information in Limit Cycle Oscillators. Physical Review Letters. 2009 Dec;103(24):1–4. Available from: http://link.aps.org/doi/10.1103/PhysRevLett.103.248105.

54 Mato G, Samengo I. Type I and type II neuron models are selectively driven by differential stimulus features. Neural Computation. 2008 Oct;20(10):2418–40. Available from: http://www.ncbi.nlm.nih.gov/pubmed/18439139.

55 Honeycutt R. Stochastic Runge-Kutta algorithms. II. colored noise. Physical Review A. 1992;45(2). Available from: http://adsabs.harvard.edu/abs/1992PhRvA.45.604H.

56 Rice SO. Mathematical analysis of random noise. Bell System Technical Journal. 1944;23(3):282–332.

57 Jung P. Threshold devices: Fractal noise and neural talk. Physical Review E. 1994;50(4):2513–2522.

58 Brette R. What Is the Most Realistic Single-Compartment Model of Spike Initiation? PLOS Computational Biology. 2015;11(4):e1004114. Available from: http://dx.plos.org/10.1371/journal.pcbi.1004114.

59 Hille B. Ion Channels of Excitable Membranes; 2001.

60 Llinás R, Yarom Y. Oscillatory properties of guinea-pig inferior olivary neurones and their pharmacological modulation: an in vitro study. Journal of physiology. 1986;376:163–182.

61 Lampl I, Yarom Y. Subthreshold oscillations of the membrane potential: a functional synchronizing and timing device. Journal of Neurophysiology. 1993 Nov;70(5):2181–6. Available from: http://www.ncbi.nlm.nih.gov/pubmed/8294979.

62 Hutcheon B, Miura RM, Yarom Y, Puil E. Low-threshold calcium current and resonance in thalamic neurons: a model of frequency preference. Journal of Neurophysiology. 1994 Feb;71(2):583–594. Available from: http://jn.physiology.org/content/71/2/583.abstract.

63 Leung LS, Yu HW. Theta-frequency resonance in hippocampal CA1 neurons in vitro demonstrated by sinusoidal current injection. Journal of Neurophysiology. 1998;79(3):1592–1596.

64 Gutfreund Y, Yarom Y, Segev I. Subthreshold oscillations and resonant frequency in guinea-pig cortical neurons: physiology and modelling. Journal of Physiology. 1995;483 ( Pt 3: 621–640.

65 Tateno T, Harsch A, Robinson HPC. Threshold firing frequency-current relationships of neurons in rat somatosensory cortex: type 1 and type 2 dynamics. Journal of Neurophysiology. 2004 Oct;92(4):2283–94. Available from: http://www.ncbi.nlm.nih.gov/pubmed/15381746.

66 Engel Ta, Schimansky-Geier L, Herz AVM, Schreiber S, Erchova I. Subthreshold membrane-potential resonances shape spike-train patterns in the entorhinal cortex. Journal of Neurophysiology. 2008 Sep;100(3):1576–89. Available from: http://www.pubmedcentral.nih.gov/articlerender.fcgi?artid=2544463&tool=pmcentrez&rendertype=abstract.

67 Muresan RC, Savin C. Resonance or integration? Self-sustained dynamics and excitability of neural microcircuits. Journal of Neurophysiology. 2007 Mar;97(3):1911–30. Available from: http://www.ncbi.nlm.nih.gov/pubmed/17135469.

68 Hutt A, Buhry L. Study of GABAergic extra-synaptic tonic inhibition in single neurons and neural populations by traversing neural scales: application to propofol-induced anaesthesia. Journal of Computational Neuroscience. 2014;37(3):417–437. Available from: http://link.springer.com/10.1007/s10827-014-0512-x.

69 Badel L, Gerstner W, Richardson MJE. Spike-triggered averages for passive and resonant neurons receiving filtered excitatory and inhibitory synaptic drive. Physical Review E - Statistical, Nonlinear, and Soft Matter Physics. 2008;78(1):1–12.

70 Pernice V, Staude B, Cardanobile S, Rotter S. How structure determines correlations in neuronal networks. PLoS Computational Biology. 2011;7(5).

71 Brunel N, Brunel N. Dynamics of sparsely connected networls of excitatory and inhibitory neurons. Computational Neuroscience. 2000;8:183–208.

72 Van Vreeswijk C, Sompolinsky H. Chaotic balanced state in a model of cortical circuits. Neural Computation. 1998 Aug;10(6):1321–71. Available from: http://www.ncbi.nlm.nih.gov/pubmed/9698348.

73 Jadi MP, Sejnowski TJ. Regulating Cortical Oscillations in an Inhibition-Stabilized Network. Proceedings of the IEEE. 2014;102(5):830–842. Available from: http://ieeexplore.ieee.org/lpdocs/epic03/wrapper.htm?arnumber=6803063.

74 Balu R, Strowbridge BW. Opposing inward and outward conductances regulate rebound discharges in olfactory mitral cells. Journal of Neurophysiology. 2007 Mar;97(3):1959–68. Available from: http://www.ncbi.nlm.nih.gov/pubmed/17151219.

75 Galan RF, Ermentrout B, Urban N. Efficient Estimation of Phase-Resetting Curves in Real Neurons and its Significance for Neural-Network Modeling. Physical Review Letters. 2005 Apr;94(15):1–4. Available from: http://link.aps.org/doi/10.1103/PhysRevLett.94.158101.

76 Bathellier B, Lagier S, Faure P, Lledo PM. Circuit properties generating gamma oscillations in a network model of the olfactory bulb. Journal of Neurophysiology. 2006 Apr;95(4):2678–91. Available from: http://www.ncbi.nlm.nih.gov/pubmed/16381804.

77 Brea JN, Kay LM, Kopell N. Biophysical model for gamma rhythms in the olfactory bulb via subthreshold oscillations. Proceedings of the National Academy of Sciences. 2009 Dec;106(51):21954–9. Available from: http://www.pubmedcentral.nih.gov/articlerender.fcgi?artid=2799880&tool=pmcentrez&rendertype=abstract.

78 Baroni F, Burkitt AN, Grayden DB. Interplay of Intrinsic and Synaptic Conductances in the Generation of High-Frequency Oscillations in Interneuronal Networks with Irregular Spiking. PLoS Computational Biology. 2014;10(5).

79 Moca VV, Nikolić D, Singer W, Muresan RC. Membrane resonance enables stable and robust gamma oscillations. Cerebral Cortex. 2014;24(1):119–142.

80 Shriki O, Hansel D, Sompolinsky H. Rate models for conductance-based cortical neuronal networks. Neural Computation. 2003 Aug;15(8):1809–41. Available from: http://www.ncbi.nlm.nih.gov/pubmed/14511514.

81 Azouz R, Gray CM. Dynamic spike threshold reveals a mechanism for synaptic coincidence detection in cortical neurons in vivo. Proceedings of the National Academy of Sciences. 2000;97(14):8110–8115.

82 Tuckwell HC. Introduction to Theoretical Neurobiology vols. 1 and 2. Cambridge University Press; 1988.

83 Renart A, Brunel N, Wang XJ. Chapter 15 Mean-Field Theory of Irregularly Spiking Neuronal Populations and Working Memory in Recurrent Cortical Networks. In: Computational Neuroscience A Comprehensive Approach; 2004. p. 431–490. Available from: http://nba.uth.tmc.edu/homepage/cnjclub/2007summer/renart_chapter.pdf.

84 Richardson MJE, Gerstner W. Synaptic shot noise and conductance fluctuations affect the membrane voltage with equal significance. Neural Computation. 2005 Apr;17(4):923–47. Available from: http://www.ncbi.nlm.nih.gov/pubmed/15829095.

85 Johannesma P. Diffusion models for the stochastic activity of neurons. Neural Networks. 1968;Available from: http://link.springer.com/content/pdf/10.1007/978-3-642-87596-0_11.pdf.

86 Monteforte M. Chaotic Dynamics in Networks of Spiking Neurons in the Balanced State [PhD thesis]. Georg-August University; 2011.

87 Brette R, Gerstner W. Adaptive exponential integrate-and-fire model as an effective description of neuronal activity. Journal of Neurophysiology. 2005 Nov;94(5):3637–42. Available from: http://www.ncbi.nlm.nih.gov/pubmed/16014787.

